# Cell type-specific effects of age and sex on human cortical neurons

**DOI:** 10.1101/2023.11.11.566717

**Authors:** Jo-Fan Chien, Hanqing Liu, Bang-An Wang, Chongyuan Luo, Anna Bartlett, Rosa Castanon, Nicholas D. Johnson, Joseph R. Nery, Julia Osteen, Junhao Li, Jordan Altshul, Mia Kenworthy, Cynthia Valadon, Michelle Liem, Naomi Claffey, Caz O’Connor, Luise A Seeker, Joseph R. Ecker, M. Margarita Behrens, Eran A. Mukamel

## Abstract

**Excitatory and inhibitory neurons establish specialized identities early in life through cell type-specific patterns of epigenetic regulation and gene expression. Although cell types are largely stable throughout the lifespan, altered transcriptional and epigenetic regulation may contribute to cognitive changes with advanced age. Using single-nucleus multiomic DNA methylation and transcriptome sequencing (snmCT-seq) in frontal cortex samples from young adult and aged donors, we found widespread age- and sex-related variability in specific neuronal cell types. The proportion of GABAergic inhibitory cells, including SST and VIP expressing cells, was reduced in aged donors. On the other hand, excitatory neurons had more profound age-related changes in their gene expression and DNA methylation compared with inhibitory cells. Hundreds of genes involved in synaptic activity were downregulated, while genes located in subtelomeric regions were upregulated with age and anti-correlated with telomere length. We further mapped sex differences in autosomal gene expression and escape from X-inactivation in specific neuron types. Multiomic single-nucleus epigenomes and transcriptomes provide new insight into the effects of age and sex on human neurons.**

## Introduction

Cell atlases have documented the diverse molecular identity and regulation of brain cell types^1^, but the impact of inter-individual differences on epigenetic and transcriptomic regulation remains unclear^2^. Epigenetic and transcriptomic signatures of neuronal identity are established prenatally and refined through childhood and adolescence, and they regulate neural function throughout the lifespan^3–5^. Altered gene expression in specific brain cell populations could contribute to age-related changes in cognition or risk for neurodegenerative disease^6,7^. Whereas single-cell transcriptome sequencing offers a snapshot of one moment in the life of a cell, epigenetic modifications such as DNA methylation represent stable and persistent signatures of brain cell regulation across cell types and individuals. Single-nucleus multi-omic sequencing of neurons from donors across the adult lifespan provides a comprehensive view of stable and age-variable cell regulation in the human brain.

## Results

### A multiomic atlas of human frontal cortex neurons across age and sex

We measured the transcriptome and DNA methylome^8^ in frontal cortex neurons of young (23-30 years old) and aged (70-74 years old) male and female donors (n=11, Fig. 1A). We focus on the human dorsolateral prefrontal cortex (Brodmann area 46, BA46), a critical region involved in cognitive control and executive function and implicated in neuropsychiatric disorders. We obtained 55,447 high-quality nuclei with an average of >1.37 million DNA reads, which were enriched for neurons (NeuN+). Most of these cells (39,830) passed stringent quality control criteria for RNA-seq data, providing on average 213,000 RNA reads, and 6,800 genes detected per cell (Fig. S1, Supplementary Table 1).

**Fig 1.**
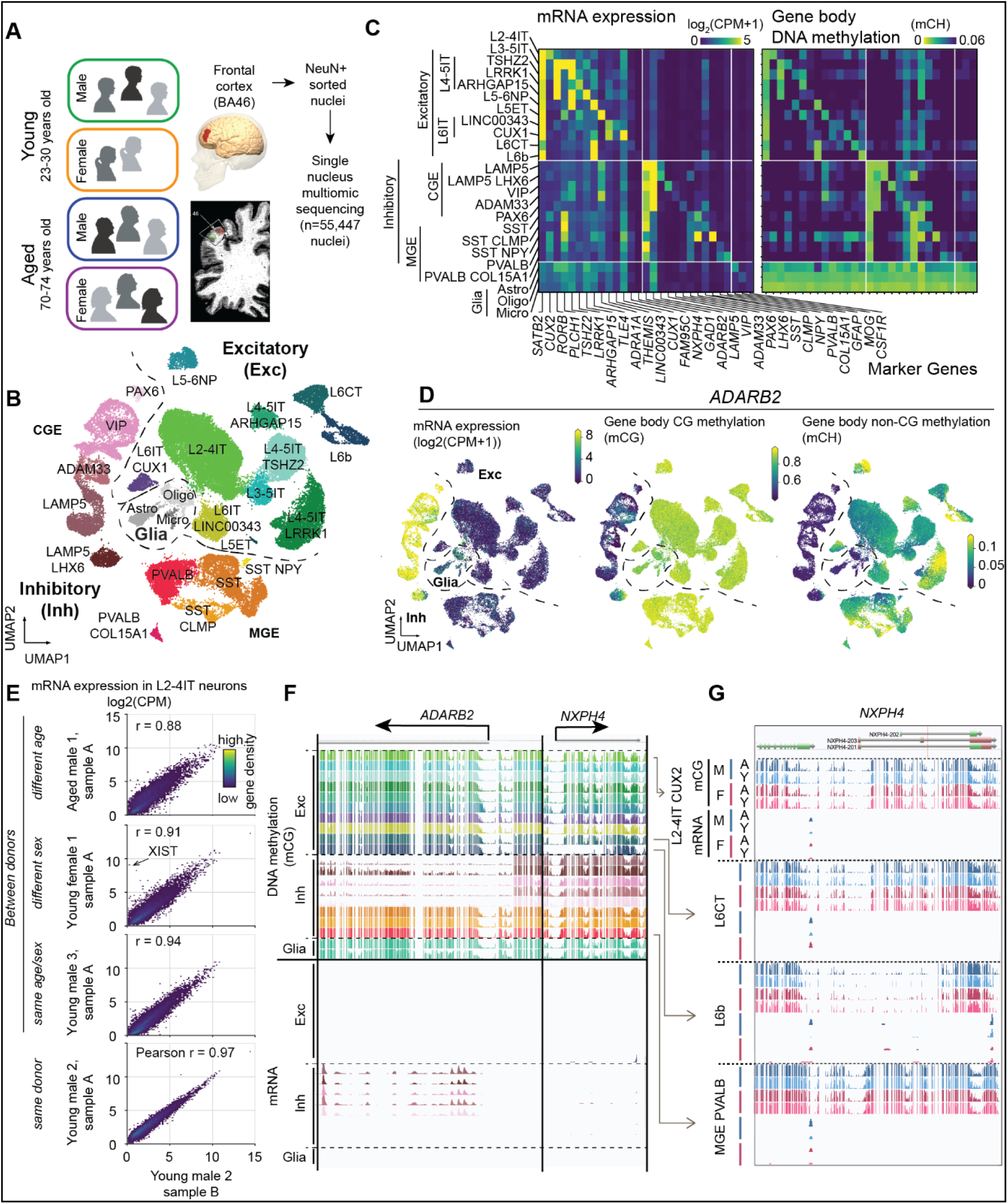
Single nucleus multiomic analysis of inter-individual transcriptomic-epigenomic diversity in human frontal cortex. (A) Study design. (B) UMAP embedding of nuclei assigned to 11 excitatory, 10 inhibitory and 3 glial types using non-CG DNA methylation (mCH) in 100 kb bins. (C) Cell type marker RNA expression and gene body mCH. (D) Correspondence of mRNA, mCG and mCH for the CGE interneuron marker, *ADARB2*. (E) Correlation of mRNA expression between independent samples from the same donor, between different donors with the same age/sex and different age or sex. (F) Pseudobulk expression (mRNA) and DNA methylation (mCG) at *ADARB2* and *NXPH4* (L6b excitatory neurons), averaged across subjects. (G) Separate tracks for young, aged, male and female groups showing consistent epigenomic and transcriptomic cell type signatures.

To assess donor variability, we annotated cells according to major neuron types rather than fine-grained subtypes. We identified 11 glutamatergic excitatory and 10 GABAergic inhibitory neuron types based on 100 kb bin non-CG methylation (mCH) features^9^, which we labeled based on expression of canonical mRNA cell-type markers^10^ (Fig. 1B,C). We also obtained a smaller number of glial cells, including astrocytes, oligodendrocytes and microglia, from some samples. The pseudobulk gene expression profiles from each cell type were highly consistent with reference cell types from scRNA-seq of multiple human cortical regions^10^ (mean Spearman correlation = 0.84 ± 0.05 for 1000 highly variable genes between cell types, Fig. S2). As expected, cell type marker gene expression correlated with low gene body DNA methylation in both CG and non-CG contexts (Fig. 1C, D, F, Fig. S3B)^8^.

RNA expression and DNA methylation features were highly consistent between independent tissue samples from the same donor. For example, samples of L2-4 intra-telencephalic (IT) cells from the same donor had Pearson correlation r = 0.98 ± 0.01 for RNA, 0.99 ± 0.01 for mCG in gene bodies, and 0.98 ± 0.01 for mCH (Fig. 1E and Fig. S3C). By contrast, the correlation of these signatures in different individuals is lower, especially between donors with different age or sex (r = 0.90 ± 0.03 for RNA, 0.97 ± 0.01 for gene body mCG, 0.91 ± 0.03 for gene body mCH, Fig. 1E and Fig. S3C). The correlation between signatures for different cell types was far lower (r = 0.73 ± 0.08 for RNA, 0.80 ± 0.07 for mCG in gene bodies, and 0.48 ± 0.15 for mCH, Fig. S3D).

These multi-omic data are displayed in a genome browser that compares base resolution DNA methylation with gene expression for each demographic group (brainome.ucsd.edu/HumanBrainAging). For example, the gene *ADARB2* is highly expressed across CGE-derived inhibitory neuron types (VIP, LAMP5) and shows corresponding patterns of low mCG and mCH across the gene body, while *NXPH4* labels a group of deep layer excitatory neurons (L6 cortico-thalamic, CT; Fig. 1E). Stratifying the genomic data by donor age and sex shows the high reproducibility of DNA methylation and RNA expression tracks for young, aged, male and female donors (Fig. 1F). These integrated multi-omic, multi-donor data allowed us to investigate age- and sex-related variability in brain cell transcriptomes and epigenomes.

### SST and VIP expressing GABAergic cells are reduced in aged frontal cortex

Cortical glutamatergic excitatory and GABAergic inhibitory neurons are produced before birth, and their finely balanced interaction throughout the lifespan regulates neural network activity to enable cognitive information processing. Although new neurons are not produced in the adult cortex, whether the ratio of excitatory to inhibitory neurons changes with age remains unclear. We found a 68% lower proportion of inhibitory neurons in aged samples (mean 0.28 ± 0.06 s.d.) compared with young samples (0.41 ± 0.06, p=0.01, Wilcoxon signed-rank test, Fig. 2A). These differences were not explained by sample quality, as we found no difference in the fraction of high-quality cells or in DNA methylation quality-control metrics in young vs. aged donors (Fig. S1).

**Fig 2.**
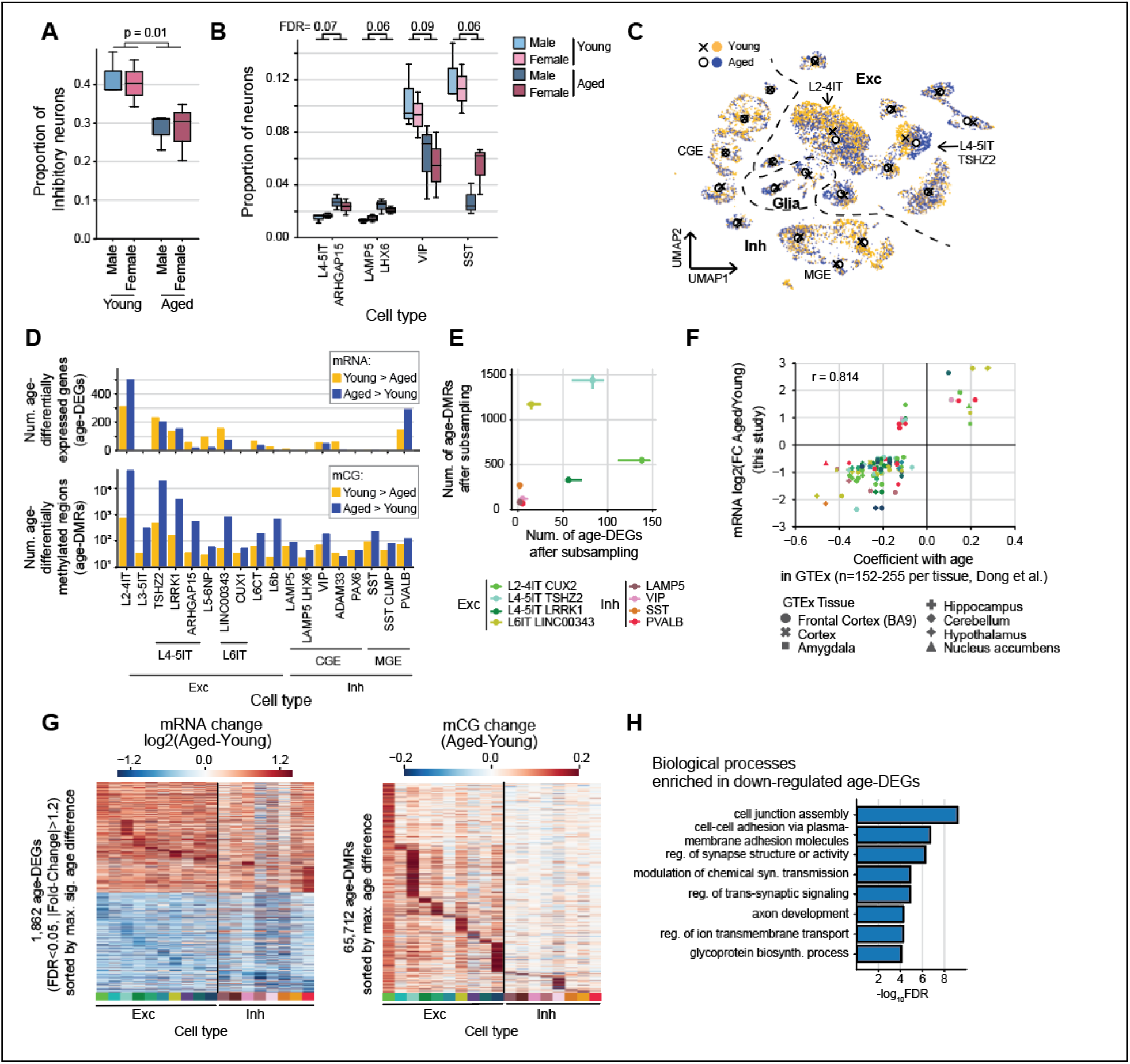
Aging effects on inhibitory and excitatory neuron cell types. (A, B) Proportion of GABAergic inhibitory neurons (A) and neuron subtypes (B) in young and aged donors (Wilcoxon rank-sum test; n=5 young, n=6 aged donors). (C) Excitatory neurons separate by age in a UMAP embedding. Markers show centroids of cells by type from young (x) and aged (circles) donors. (D) Number of age-differentially expressed genes (age-DEGs) and differentially methylated regions (age-DMRs) in each cell type. (E) Number of age-DEGs and age-DMRs after sampling 100 nuclei per cell type per sample. Error bars show the 25th and 75th quantile across random samples. (F) Correlation of age effect on gene expression in our data with GTEx data from bulk frontal cortex^17^. (G) Fold change of significant age-DEGs in at least one cell type (left), and mCG difference between aged and young neurons in age-DMRs (right). (H) Biological pathways enriched for age-DEGs that are downregulated in aged neurons (FDR <1e-3).

The age-associated decline of inhibitory neurons was driven by substantial loss of SST- (2.90-fold, FDR<0.1) and VIP-expressing cells (1.71-fold) (Fig. 2B; Fig. S4A). There was also an increase in the proportion of CGE-derived LAMP5 LHX6 neurons (1.67-fold), and a subtype of L4-5 IT excitatory neurons (1.50-fold). Notably, an increased excitatory to inhibitory neuron ratio and loss of SST or VIP neurons has been observed in the hippocampus of Alzheimer’s disease patients^11–14^.

### Widespread age-related changes in cell type-specific gene expression and DNA methylation

The deep epigenomic and transcriptomic profiles of our snmCT-seq data allowed us to further examine age-related changes in gene regulation in specific neuron types. The low-dimensional embedding of cells based on CG and non-CG DNA methylation throughout the genome showed substantial differences between excitatory neurons from young and aged donors (Fig. 2C). By contrast, inhibitory cells from all donors had very similar global methylation profiles and their UMAP embeddings were largely overlapping. We did not analyze age-related changes in glial cell types because the number of glial cells was not sufficiently balanced across donors.

The genome-wide average level of CG DNA methylation varies across neuron types^15,16^, ranging from a low level of methylation in L2-4 IT neurons (mCG=0.82) to the highest levels in SST inhibitory cells (mCG=0.86) (Fig. S4B). Non-CG methylation, particularly at CA dinucleotides, varied by more than 2-fold across neuron types, with mCA ranging from 0.07 to 0.15 (Fig. S4C, D). Genome-wide average methylation for each cell type was largely stable in aging, with slight increases in mCG in some excitatory neurons (Fig. S4B, C, D). This shows that any age-related global changes in mCG and mCH are small relative to the cell type-specific patterns of DNA methylation that are established in early life and largely persist through the adult lifespan.

We found 1,904 unique differentially expressed genes (age-DEGs) in young vs. aged neurons (FDR<0.05 and fold-change >1.2 in at least one cell type, Fig. 2D, Table S3). Overall, we found more age-DEGs in excitatory neurons (1,536 unique genes) compared with inhibitory neurons (601). In particular, there were relatively few age-DEGs in SST- (7 age-DEGs) or VIP-expression (109) inhibitory cells. This indicates that, despite their reduced population in aged donor brains, the surviving SST and VIP cells maintain a similar gene expression profile compared with young adults.

Despite the limited number of donors in our study, the aging changes were highly consistent with mRNA measurements from bulk brain tissue in a large cohort of up to n=255 donors from the gene tissue expression project (GTEx) (r = 0.71, Fig. 2F; Fig. S4D)^17^. Whereas the GTEx data identified just 28 DEGs, our single nucleus data identified more alterations to brain gene expression.

Changes in gene expression during aging could be driven by alterations to epigenetic modifications, including DNA methylation. Thousands of cytosines throughout the genome have systematic changes in DNA methylation during childhood brain development^3,4^, and the DNA methylome provides a clock-like signature of aging throughout the lifespan in multiple tissues and species^18,19^. However, previous analyses of age-related changes in DNA methylation lacked cell type-specific resolution, and did not assess the methylation status of all CG and non-CG sites genome-wide. Focusing on mCG, we called differentially methylated regions (DMRs) between aged and young donors in each neuron type, controlling for sex^20,21^. We found age-DMRs in all cell types, with more regions gaining than losing methylation with age (Fig. 2D, Table S4). For example, we found 19,715 age-DMRs in L4-5 IT-TSHZ2 neurons (covering 8.5 Mbp, ∼0.3% of the genome), and 98% of these DMRs increase methylation in aged samples (hypermethylated age-DMRs).

To directly compare age-related changes in expression across cell types without bias due to differences in cell abundance, we sampled 50 cells from each of the major neuron types for each sample followed by DE gene analysis (Fig. 2E). We also applied Augur to prioritize cell types that have the most age effect^22^ (Fig. S4E). Although the number of significant DEGs and DMRs was greatest in excitatory IT neurons, the effect of aging was similar across neuron types for most age-DEGs even in cell types in which the effect did not reach statistical significance (Fig. 2G). Among the 1,904 age-DEGs, only two protein-coding genes had opposite direction of age-related change between excitatory and inhibitory neurons (*ATP2B4*: 3.3-fold increase in L4-5IT TSHZ2 and 2.5-fold decrease in MGE PVALB; *INPP4B*: 3.8-fold increase in L6CT and 2.2-fold decrease in CGE ADAM33). By contrast, age-DMRs are more cell type specific and more prominent in excitatory neurons, especially in L2-4 IT and L4-5 IT TSHZ2 (Fig. 2G).

### Reduced expression and increased methylation of synaptic genes in the aging brain

Genes down-regulated with age are enriched in pathways related to synaptic signaling and cell adhesion (Fig. 2H). These included neuroligin 1, *NLGN1*, which mediates the formation and maintenance of excitatory synapses^23^; phosphatase and actin regulator 1, *PHACTR1*, encoding a synaptic protein regulating excitatory synapses^24^; and the voltage-gated potassium channel *KCNH7^25^*(Fig. S5D). The downregulation of synaptic and cell adhesion genes may be associated with reduced synaptic plasticity^26,27^. By contrast, only a single ontology category, histone H4 acetylation, was significantly enriched in up-regulated genes in inhibitory neurons (FDR < 0.01, Fig. S5C) (see below).

### Increased DNA methylation in aged neurons correlates with reduced gene expression

DNA methylation differences between cell types correlates with cell type-specific expression^15,28^, but whether age-related changes to the epigenome correlate with transcriptomic age changes has not been established. To address this, we calculated the Spearman correlation of the fold-change (aged/young) for RNA vs. DNA methylation features at age-DE genes (Fig. 3A, Fig. S6). Gene body mCH and mCG had a strong negative association with mRNA expression in most neuron types (FDR < 0.01). For example, excitatory L4-5 IT RORB TSHZ2 had r = -0.53 (p<10^-31^, Fig. 3B). Gene body mCG had a similar correlation, but mCG and mCH at the promoter region (transcription start site ± 1kb) was less strongly correlated (Fig3A, Fig. S6). Correlating mRNA with mC features across all genes showed a similar pattern of negative association (Fig. S6C). We conclude that changes in neuronal DNA methylation coincide with age-related expression changes in the adult brain, consistent with a potential regulatory role.

**Fig 3.**
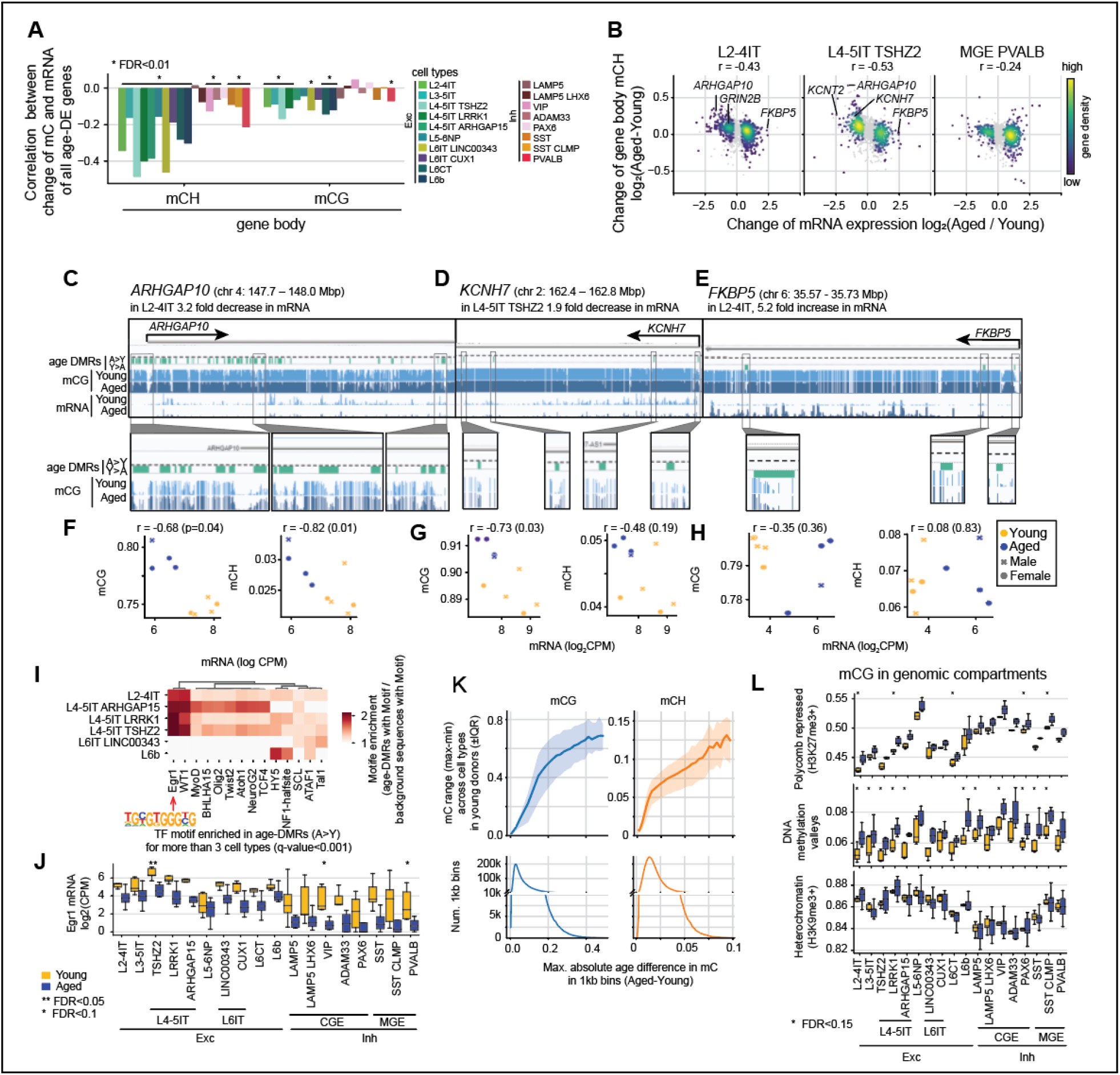
Correlated transcriptome and DNA methylome changes in the aged human brain. (A) Spearman correlation coefficient between the age change of gene body DNA methylation (aged - young) and the fold-change in gene expression in age-DEGs.* FDR <0.01. (B) Example scatter plots showing significant age-DEGs (colored points) correlated with the age change in gene body mCH. Gray points show all age-DEGs in at least one cell type. (C,D) Browser view of the *ARHGAP10* (C) and *KCNH7* (D) loci, showing reduced gene expression and increased mCG at age-DMRs throughout the gene body. (E) Browser view of the *FKBP5* loci, showing reduced mCG at age-DMRs and increased gene expression. (F,G,H) Scatter plots of mRNA expression and CG, non-CG methylation for each donor in *ARHGAP10* (F), *KCNH7* (G) and *FKBP5* (H). (I) Motif enrichment of hypermethylated age-DMRs. Only motifs significantly enriched (FDR<0.001) in more than two cell types are shown. (J) Gene expression of *EGR1* gene. (K) The relation between maximum methylation difference in age and the methylation difference between cell types for all 1kb bins. Bottom row is the histogram of the bins with age methylation difference. (L) mCG age difference in genomic compartments. Polycomb and heterochromatin regions are called by chromHMM whole brain tissue annotation. DNA methylation valleys are called by methylSeekR from our data.

To illustrate the age-related changes in gene expression and DNA methylation, we highlight the rho GTPase activating protein 10, *ARHGAP10* (Fig. 3C). mRNA for *ARHGAP10* is highly expressed in upper layer excitatory neurons (L2-4 IT) in young donors (>435 transcripts per million mapped reads, TPM), and subsequently reduced 3.2-fold in older donors. Consistent with this downregulation, we found several DMRs around the *ARHGAP10* locus where DNA methylation increases with age in L2-4 IT neurons. DMRs occurred in the upstream intergenic region, marking putative cis-regulatory elements, as well as within the promoter and gene body. Gene body mC also had a strong negative correlation with *ARHGAP10* mRNA expression across donors (r = –0.68 for mCG, –0.82 for mCH, Fig. 3F). Similarly, the highly expressed potassium voltage-gated channel subfamily H member 7, *KCNH7*, was reduced 1.9-fold in aged donors and showed a corresponding increase of CG methylation (Fig. 3D, G).

In total, age-DMRs covered 32.4 Mbp, mainly at non-exonic sequences (94%). The majority of the age-DMRs were in introns (69%) or intergenic regions (25%). To further characterize the DMRs, we analyzed their enrichment in functional genomic compartments defined by multiple histone modification ChIP-seq datasets from human prefrontal cortex using chromHMM^29^ (Fig. S5F). DMRs that increase methylation in aged neurons were enriched in enhancers, and in states associated with Polycomb repressive complex 2 (PRC2). DMRs that lose methylation in aged neurons were only enriched in active states.

To explore the significance of DNA methylation changes we used sequence motif enrichment analysis to identify transcription factors that may bind at age-DMRs^30^. Motifs of early growth response (EGR) family factors, including WT1 and EGR1, were enriched in excitatory neuron age-DMRs that increase methylation with age (FDR<0.001, Fig. 3I). Up to 12.5% of hypermethylated (aged > young) age-DMRs overlapped EGR1 motifs (Fig. S5G). *EGR1* expression was also strongly reduced with age (4-30 fold) in both excitatory and inhibitory neurons (Fig. 3J), although there was no significant difference in gene body methylation. *EGR1* is an immediate early gene encoding a zinc finger transcription factor that is important for long-term memory consolidation^31,32^. Our findings in the human frontal cortex extend previous observations of reduced *EGR1* expression and increased promoter mCG in the hippocampus of aged mice^33^. By contrast, regions that lost mCG in aged neurons were enriched in transcription factor binding motifs of the AP-1 family of immediate early genes, Fos and Jun, and the transcription elongation factor GRE (Fig. S5H).

DNA methylation plays a fundamental role in development, establishing unique sets of hypomethylated DMRs that mark each mature cell type^9,16^. These regions have large differences in mCG across adult neuron types. We hypothesized that age-related changes in DNA methylation may specifically affect these developmentally programmed sites of cell type regulation. Consistent with this, we found that regions with the greatest difference in mCG or mCH between aged and young cells in any cell type also had large differences in mean methylation between cell types in young adults (Fig. 3K).

### Low methylated regions gain mCG with age

Studies of the DNA methylation clock have identified systematic changes in mCG throughout the lifespan, particularly at sites with low mCG^34^. Consistent with this, we found increased mCG with age in DNA methylation valleys (DMVs), defined as extended regions (≥5 kb) with low average mCG (<15%)^35^ (Fig. 3L). DNA methylation also increased at regions associated with Polycomb repressive complex 2 (PRC2) in prefrontal cortex^29^, although the PRC2 annotation was not cell type-specific. By contrast, there was no effect of age on mCG at heterochromatin regions marked by the repressive histone modification H3K9me3 (Fig. 3L). The lack of increased mCG in heterochromatin regions is notable given the observation of increased chromatin accessibility and reduced H3K9me3 at these sites in excitatory neurons in aged mice^36^.

### Increased expression of subtelomeric genes in aged brain cells

We observed notable examples of genes with strongly increased expression in aged neurons. *FKBP5*, a gene involved in immunoregulation and glucocorticoid receptor signaling, increased expression 5.2-fold in aged neurons and harbored hypomethylated (young > aged) age-DMRs in its gene body (Fig. 3E). Aging and stress have been associated with increased *FKBP5* expression and decreased methylation at two CG sites in blood in multiple large human cohorts (n>3000), and this was mechanistically linked to NF-k-B driven inflammation^37,38^. Another example of an age-increased gene in L2-4 IT neurons was the mu opioid receptor, *OPRM1* (FDR<0.05, >2-fold increase). Altered opioid receptor signaling has been reported in aged rodents^39^.

Despite these examples, we did not identify a clear functional association for age-increased genes as a group, in contrast with the significant enrichment of synaptic function for age-decreased genes (Fig. 2H). Moreover, whereas age-decreased genes fit an expected pattern with increased methylation and enriched hypermethylated age-DMRs in their gene body, age-increased genes were not enriched in gene-body mCG (Fig. 4A, B). Instead, genomic regions that surround age-increased genes had less age-related accumulation of mCH compared with regions around age-decreased genes (Fig. 4A). mCG hypo-DMRs (aged < young) were slightly enriched at the promoters of age-increased genes (Fig. 4B).

**Fig 4.**
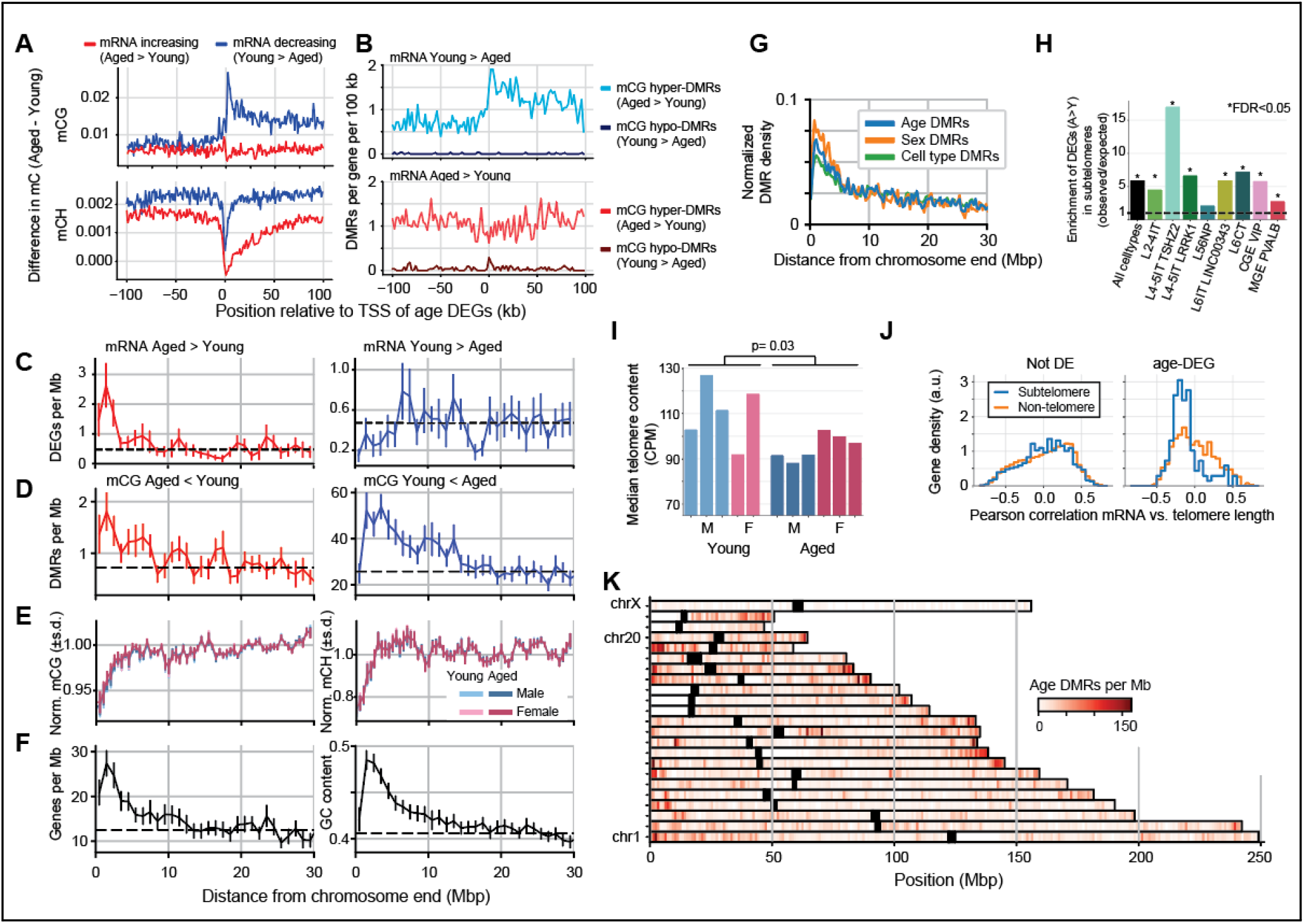
Increased expression of subtelomeric genes in aged neurons. (A) Average DNA methylation around the promoter of age-DEGs whose expression increases (red) or decreases (blue) with age. Age-decreasing genes have higher gene body mCG, while age-increasing genes have lower mCH both in the gene body and in the surrounding region. (B) Density of differentially methylated regions (DMRs) around age-DEGs. Hypermethylated age-DMRs are enriched in age-decreasing genes. Hypomethylated age-DMRs are rare, but they are enriched in the promoters of age-increasing genes. (C-F) Distribution of DEGs (C), DMRs (D), normalized DNA methylation (E), genes and genomic GC content (F) as a function of distance to the nearest chromosome end. Error bars are the standard error across chromosomes. (G) Normalized density of age-, sex- and cell type-DMRs as a function of distance to the nearest chromosome end. (H) Enrichment of age-increasing DEGs in subtelomeres (≤5 Mb from chromosome end) (FDR <0.05, Fisher exact test). Enrichment calculated as (observed/expected) subtelomeric A>Y genes, where expected = (# A>Y genes)(# subtelomeric DEGs) / (# DEGs). (I) Median telomere length in L2-4IT neurons for each donor, estimated by TelomereHunter^40^. (J) Pearson correlation of pseudobulk mRNA expression with donor telomere length for DE and non-DE genes in subtelomeric or non-telomeric regions in L2-4IT neurons. (K) Map of the density of age DMRs across the genome.

Given the reduced mCH around age-increased genes, we reasoned that these genes may be clustered in genomic regions with a distinct chromatin organization. We found that age-increased genes, but not age-decreased genes, were enriched ∼1.5-fold within the subtelomeric regions, i.e. <5 Mb from the ends of chromosomes (Fig. 4C). We found a similar pattern of enrichment for age DMRs, including both hypo- and hyper-mCG age-DMRs (Fig. 4D). The subtelomeric regions have lower average mCG (∼5% lower than the genome average) and mCH (∼20% lower, Fig. 4E). Subtelomeres also have higher gene density and higher GC content than the rest of the genome (Fig. 4F).

Telomere attrition and telomere DNA damage response in aging have been reported in both dividing and post-mitotic cells^41,42^. We reasoned that the enrichment of age-increasing DEGs in subtelomere regions might be related to the disruption of the chromatin structure at chromosome ends due to telomere damage. We applied TelomereHunter^40^ to estimate telomere length for each donor, using information from snmCT-seq reads that mapped to unique genomic locations as well as unmappable reads. Estimated telomere length was significantly shorter in L2-4 IT neurons from aged vs. young donors (Fig. 4I, p<0.05, Wilcoxon rank-sum test). Furthermore, shorter telomere length correlated with increased expression of age-DEGs located in the subtelomere regions, consistent with a telomere position effect^43^ (Fig. 4J).

We found age DMRs enriched at the ends of many, but not all, chromosomes (Fig. 4K). By contrast, there was no enrichment at centromeres. The enrichment of DMRs was not unique to age-related mC, but was also observed for sex DMRs and cell-type DMRs (Fig. 4J). This enrichment was also not unique to a particular cell type but was observed across multiple excitatory and inhibitory neuron types (Fig. 4H, FDR<0.05).

These findings are notable given the observation of age-increased expression of genes in subtelomeric regions in multiple tissues across a large cohort (GTEx)^17^. In that study, six tissues had significant enrichment of age-increased genes within the subtelomeres, including whole blood, skeletal muscle, thyroid, and sigmoid colon, all of which have been shown to progressively lose telomere length with advanced age. The GTEx data showed no enrichment of age-increased genes in the subtelomeres for any of the 14 brain tissues tested, including frontal cortex^17^. Despite the lower number of donors in our study, our single-cell data identified a larger number of age-DEGs and recapitulated the enrichment of age-increased DEGs observed in other tissues. These data indicate that post-mitotic neurons experience age-related alterations of gene expression, including cell type specific age-increased expression, in the subtelomeric compartment.

### Transcriptome and DNA methylome differences between male and female neurons

In contrast with the effect of age, sex differences in cell type proportion and global CG or CH DNA methylation were not significant (Fig. S4A,S7A,B, FDR>0.1)^2^. We detected a total of 160 autosomal and 6 X-linked differentially expressed genes (sex-DEGs) across all cell types; Y-linked genes were excluded from this analysis (Fig. 5A, Table S5). We also identified 11,702 unique autosomal sex-DMRs (merged across cell types), covering 3.1 Mb (Table S6). As previously reported, sex-DMRs also occured throughout chrX, covering 108 Mb (69% of the whole chromosome)^45,46^.

**Fig 5.**
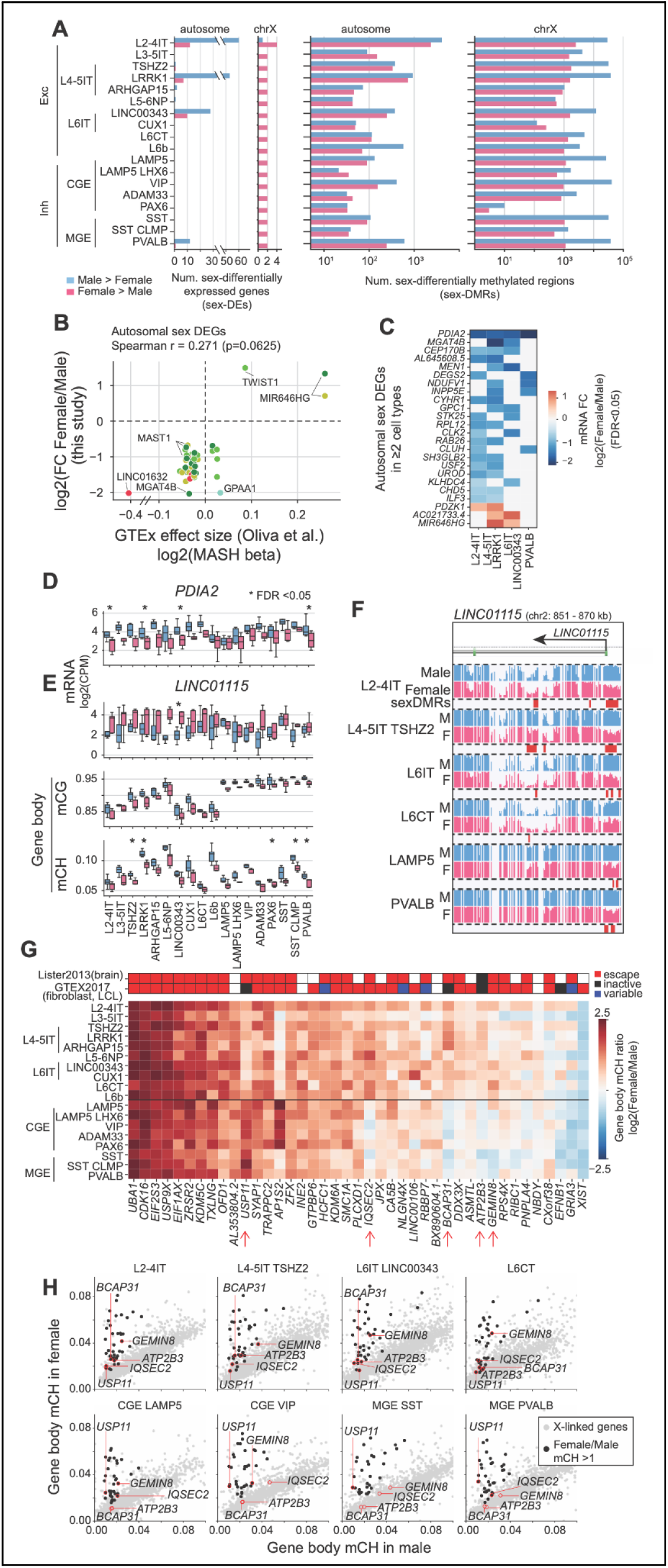
Sex differences in DNA methylation and gene expression in human neurons. (A) Number of autosomal and X-linked sex differentially expressed genes (sex-DEGs) and differentially methylated regions (sex-DMRs) in each cell type. (B) Comparison of the female/male expression for autosomal sex-DEGs in neural cell types with GTEx data from bulk frontal cortex^44^ (colors correspond to cell types shown in Fig. 3A). (C) Fold-change of autosomal sex-DEGs that are significant in multiple cell types. (D, E) Sex-specific RNA expression of *PDIA2*, and RNA and gene body DNA methylation of *LINC011115*. (F) CG DNA methylation in the promoter and 5’ intronic region of *LINC011115*, with female hypo-mCG age-DMRs. (G) Gene body non-CG methylation ratio (female/male) in 41 putative chrX escape genes with female mCH > 0.02 and female/male mCH >1.5 in at least one cell type. (H) Scatter plots of gene body mCH in male and female neurons for X-linked genes (gray dots) and putative escape genes (black). Cell type-specific X-inactivation escape genes are labeled.

Sex-specific differential expression in our dataset was consistent with analysis of whole-tissue RNA-seq in two studies using GTEx data from bulk brain tissue samples (Fig. S7C, D)^44,47^. The ratio of gene expression in female vs. male cells for autosomal sex-DEGs correlated with GTEx frontal cortex (Brodmann area 9)^44^. The strongest correlation was with L2-4 IT neurons (Spearman r=0.22, p<10^-10^), which represent the largest subpopulation of neurons in the frontal cortex (Fig. 5B, S7D). A notable example of an autosomal gene with strong sex-biased expression in our data was protein disulfide isomerase family A member 2, *PDIA2* (Fig. 5C,D). *PDIA2,* which is reported to modulate estradiol activity through direct binding^48^, is expressed 3.4-4.4 fold more in male than female neurons.

Among the thousands of autosomal sex-DMRs, there was a similar number of regions with increased methylation in males and in females. As an example, the long non-coding RNA, *LINC01115,* had a prominent pattern of mCG DMRs (female < male) at the promoter and 5’ intronic region in both excitatory and inhibitory neurons (Fig. 5F). The gene also had significantly lower gene body mCH in females than males. Consistent with this female hypo-methylation, *LINC01115* was expressed more strongly in females than males (Fig. 5E). *LINC01115* was recently reported to have a striking increase in the active chromatin mark H3K4me3 in the prefrontal cortex of schizophrenic patients compared to controls^49^.

### Cell type-specific X-inactivation escape genes

X chromosome inactivation balances gene expression on chrX in females and males. This process prevents accumulation of mCH across the majority of the female inactive X chromosome^45^. A subset of X-linked genes escapes X inactivation and accumulates high levels of mCH in the gene body in female neurons^3,45,46^. Previous studies using bulk tissue samples could not identify cell type-specific X-inactivation escape features. Using our single cell data, we identified 41 chrX genes with female-to-male gene body mCH ratio greater than 1.5 in at least one cell type, which we defined as putative escapee genes (Fig. 5F,G). Previously identified escapees^3,50^ such as *UBA1* and *KDM5C* had consistent escape signatures across all cell types. However, some genes previously identified as escapees exhibited cell type-specific non-CG methylation differences between males and females. For instance, *GEMIN8* had higher gene body non-CG methylation in females compared to males in excitatory neurons and CGE-derived inhibitory neurons, but not in MGE-derived inhibitory neurons. *IQSEC2* lacks escape signatures in VIP and SST neurons. Moreover, genes previously considered to be subject to inactivation exhibited cell type-specific escape signatures in our data. For example, *BCAP31* and *ATP2B3* had elevated female non-CG methylation in excitatory neurons but not in inhibitory neurons. By contrast, *USP11* had strong escape signatures in inhibitory neurons but not in excitatory neurons.

## Discussion

No two humans are alike, and inter-individual differences in brain cell regulation are critical to understanding the basis of human cognitive diversity. Our study adds the dimensions of age- and sex-related variation to our knowledge of the transcriptomic and epigenomic diversity of human neuron types.

Studies of the so-called DNA methylation clock have shown that changes in methylation at several dozen sites throughout the genome constitute a broadly shared predictive signature of age with similar features across human tissues and mammalian species^18,34,51^. Our data using whole-genome sequencing of the DNA methylome and transcriptome in the adult human cortex reveal cell type specific age-related changes in each of the specialized excitatory and inhibitory neuron types.

We found that intratelencephalic-projecting (IT) excitatory neurons in layers 2-4 and 4-5 were most affected by aging in terms of both gene expression and DNA methylation. Age-related changes in DNA methylation were strongly correlated with changes in gene expression. Increased DNA methylation at both CG and non-CG sites accompanied decreased gene expression, particularly for genes implicated in synaptic structure and function^26^ such as *Egr1^33^, Nlgn1^23^, PHACTR1^24^,* and *Kcnh7^25^*. Moreover, distal enhancers with age-increased CG methylation (age-DMRs) were enriched in the DNA sequence motif associated with *Egr1* transcription factor binding sites. These data extend the known role of DNA methylation in pre- and post-natal brain development, when cell type specific patterns of CG and non-CG methylation are established as neurons acquire and consolidate their mature identity^3^. Our findings show that parallel changes in the DNA methylome and transcriptome of neurons throughout the lifespan contribute to age-related alterations in neural function, with the greatest impact on upper-layer excitatory neurons.

Whereas age-decreasing genes had a clear synaptic function, we found that age-increasing genes were instead characterized by a distinctive genomic distribution. These genes were enriched in the subtelomeric regions near the ends of chromosomes, consistent with observations in other tissues in which telomeres shorten with age^17^. Moreover, we found evidence for shorter telomeres in neurons from aged compared to young donors, although telomere length estimates from our snmCT-seq data are difficult to validate^40^. These observations suggests that up-regulation of gene expression in subtelomeres of post-mitotic neurons may not be related to cell division. Instead, telomere shortening could potentially be related to DNA damage repair in neurons^52,53^. Although there was no pattern of enrichment for specific functional annotations with subtelomeric age-increased genes, these genes included some with notable roles such as the glucocorticoid receptor co-chaperone *Fkbp5^37,38^*.

In contrast with excitatory neurons, GABAergic inhibitory cells had highly similar gene expression and DNA methylation patterns in young adult and aged humans. Recent studies have noted the relative conservation of inhibitory compared to excitatory neuron gene expression across brain regions, mammalian species, and inidividual human donors^10,54,55^. Instead of altered regulation, we observed a strong reduction in the proportion of two GABAergic neuron subtypes, SST and VIP expressing cells, in aged compared with young adult donors. One of these subtypes, the SST neurons, were reportedly reduce in AD patients^7,11–14^. This observation could partly reflect differences in the efficiency for sampling nuclei from distinct neuron types using our snmCT-seq procedure, which relies on FANS sorting. More direct cell counting using microscopy and/or spatial transcriptome data are needed to further validate this observation.

Our study has several limitations. First, we performed deep sequencing of >5,000 cells from each of 11 donors, a design that enables analysis of specific brain cell types at the expense of a broader representation of variability across more donors. The consistency of our age- and sex-DEGs and of the pattern of subtelomeric enrichment of age-increasing genes with GTEx data from hundreds of donors^17,44,47^ validates the reliability of our sample. As noted above, our observation of the depletion of SST and VIP cells in aged donors could be impacted by artifacts due to nuclei sorting. Finally, our study focuses on neurons, which neglects the important role of glia in aging and age-related disease^56^.

Single-cell sequencing has revolutionized biological studies by revealing the distinct regulation of individual cells. Our study shows that multiomics can extend this powerful approach to assess the dimensions of inter-individual diversity, including age and sex. Differences in brain cell regulation between individuals are an important aspect of human brain function and must be addressed as we seek a comprehensive understanding of brain cell diversity^1^.

## Supporting information

Supplemental Table 1 - donor metadata

Supplemental Table 2 - cell counts

Supplemental Table 3 - Age-DEGs

Supplemental Table 4 - Age-DMRs

Supplemental Table 5 - Sex-DEGs

Supplemental Table 6 - Sex-DMRs

## Data availability

Data from this study are available from GEO at accessions GSE193274, GSE193296, GSE193299, GSE193313, GSE193339, GSE193372, GSE193458, GSE193499, GSE201248, GSE201830, GSE201910, GSE201933, GSE202033, GSE202062, GSE202125, GSE202162. The data may be interactively explored using a custom genome browser (https://brainome.ucsd.edu/HumanBrainAging) and a single cell browser (https://cellxgene.cziscience.com/collections/91c8e321-566f-4f9d-b89e-3a164be654d5).

## Acknowledgments

This study was supported by a grant from the Chan Zuckerberg Initiative (CZI) Seed Networks for the Human Cell Atlas (CZF2019-002457). We are grateful to Xin Jin for valuable feedback and discussions.

## Author contributions

Conceptualization, JC, JRE, MMB, EAM; Methodology, JC, HL, BAW, CL, JRE, MMB, EAM; Sequencing data production, HL, BAW, AB, RC, NDJ, JRN, JO, JA, MK, CV, ML, NC, COC; Formal analysis, JC, JL, EAM; Writing, JC, EAM; Writing - review and editing, All authors; Supervision, JRE, MMB, EAM.

## Supplemental figures

**Fig S1.**
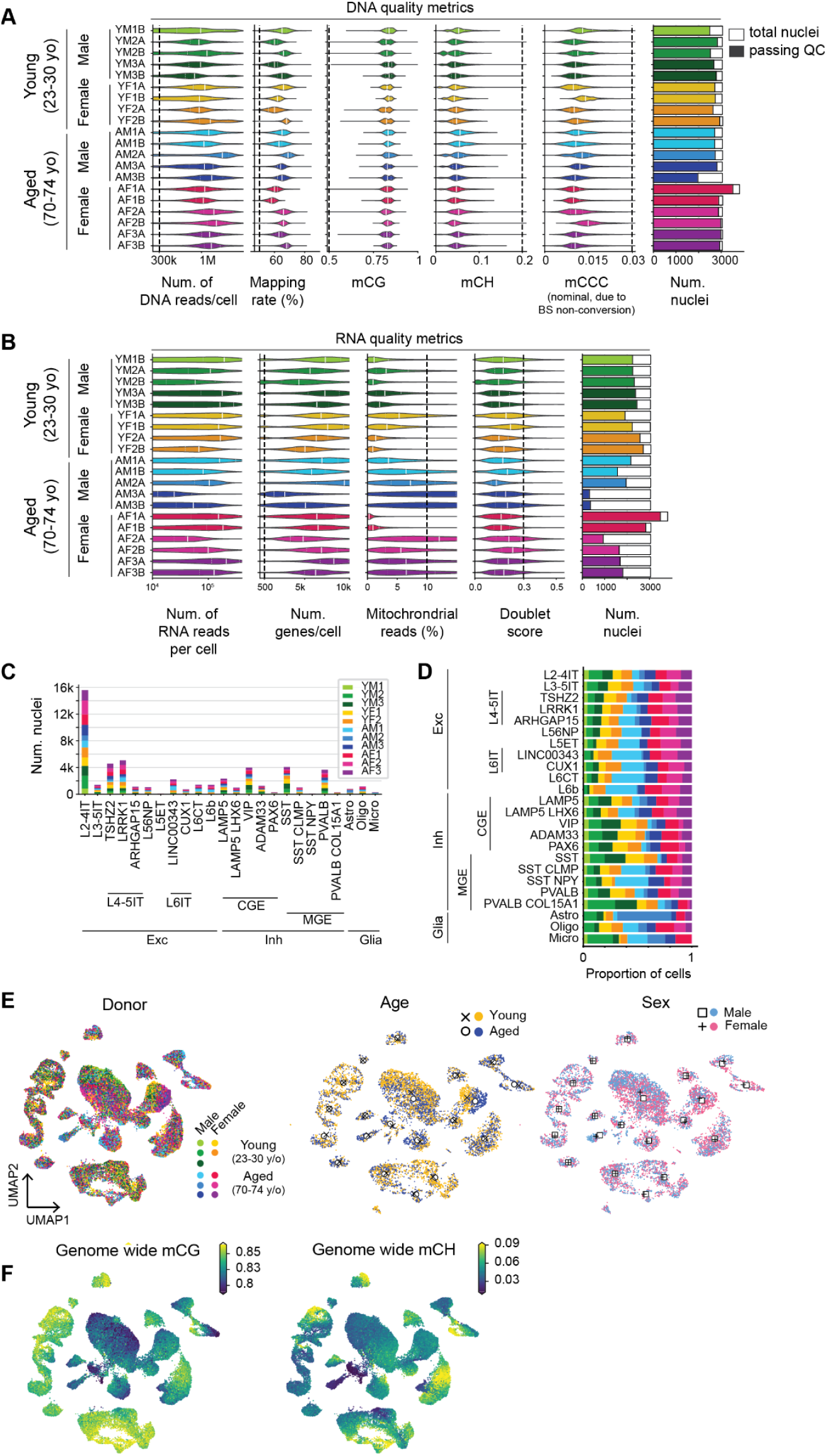
Quality control (QC) metrics for snmCT-seq data. (A) QC for DNA. Number of DNA reads, mapping rate, genome-wide mCG, mCH, mCCC level for all nuclei grouped by samples. The last column shows the number of total nuclei (open) and the number of nuclei passing DNA QC for each donor (filled, DNA unique mapped read count > 300,000, mapping rate > 0.5, mCG > 0.5, mCH level < 0.2, and mCCC < 0.05). (B) QC for RNA. Number of RNA reads, number of detected genes, percentage of reads mapped to the mitochondrial genome, and doublet score for all nuclei grouped by samples. The last column shows the total number of nuclei (open) and the number of nuclei passing RNA QC for each donor (filled, number of detected genes > 500, percent of mitochondria reads < 10%, and doublet score < 0.3). (C) Number of nuclei per cell type per donor. (D) Proportion of nuclei. (E) UMAP embedding of nuclei colored by donor, donor age, donor sex. Markers show centroids of cells by type from young (x) and aged (circles) or male (square) and female (plus) donors. (F) UMAP embedding of nuclei colored by genome-wide mCG, mCH.

**Fig S2.**
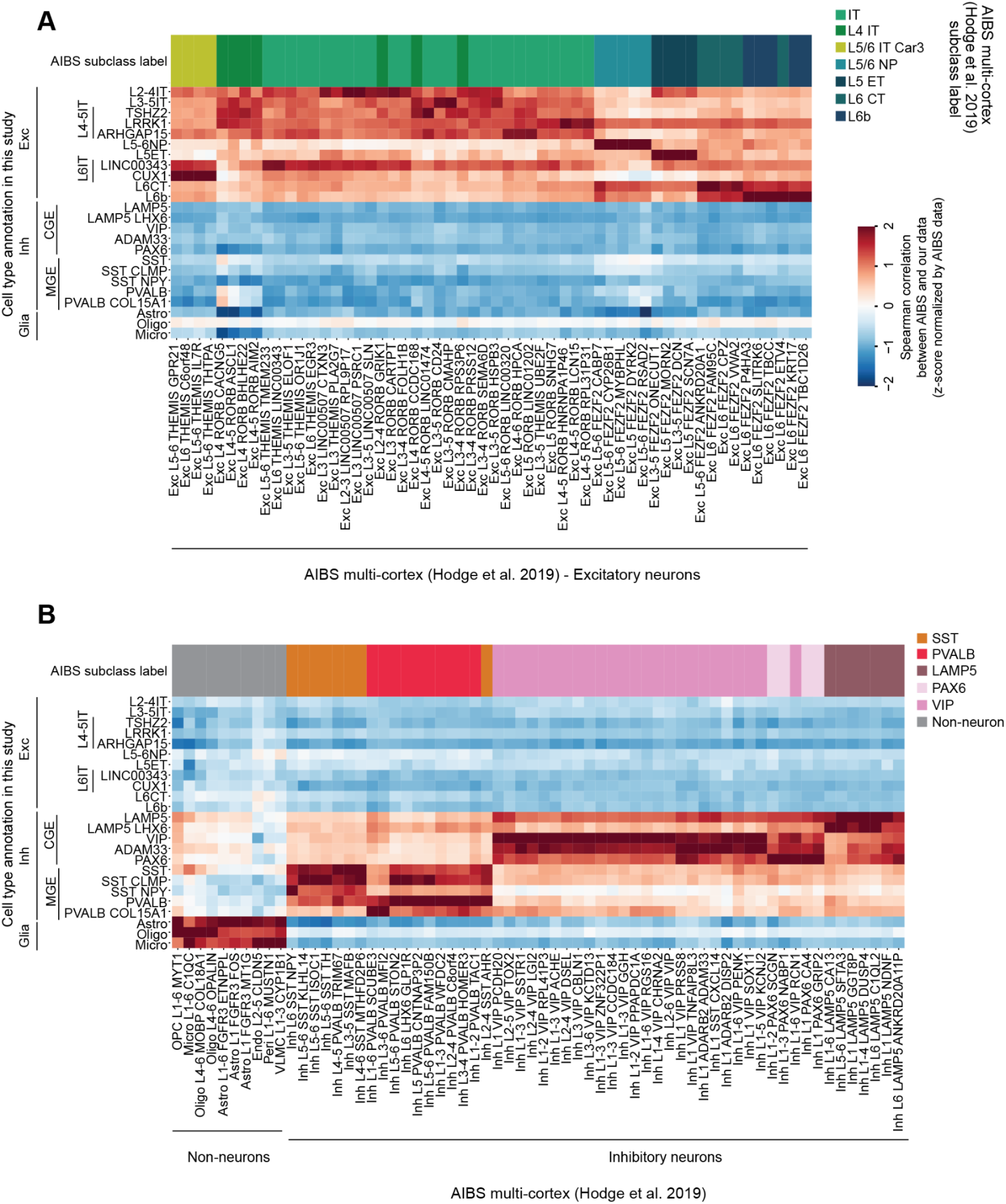
Comparison of RNA expression from snmCT-seq data (this study) with published snRNA-seq data^10^. Heatmap shows Spearman correlation between pseudobulk RNA expression (log2(CPM+1)) in our data and pseudobulk RNA expression (log2(CPM+1)) from AIBS multi-cortex data for excitatory (A) and inhibitory neurons or non-neurons (B). The correlation is calculated using the top 500 most variable autosomal protein-coding gene.

**Fig S3.**
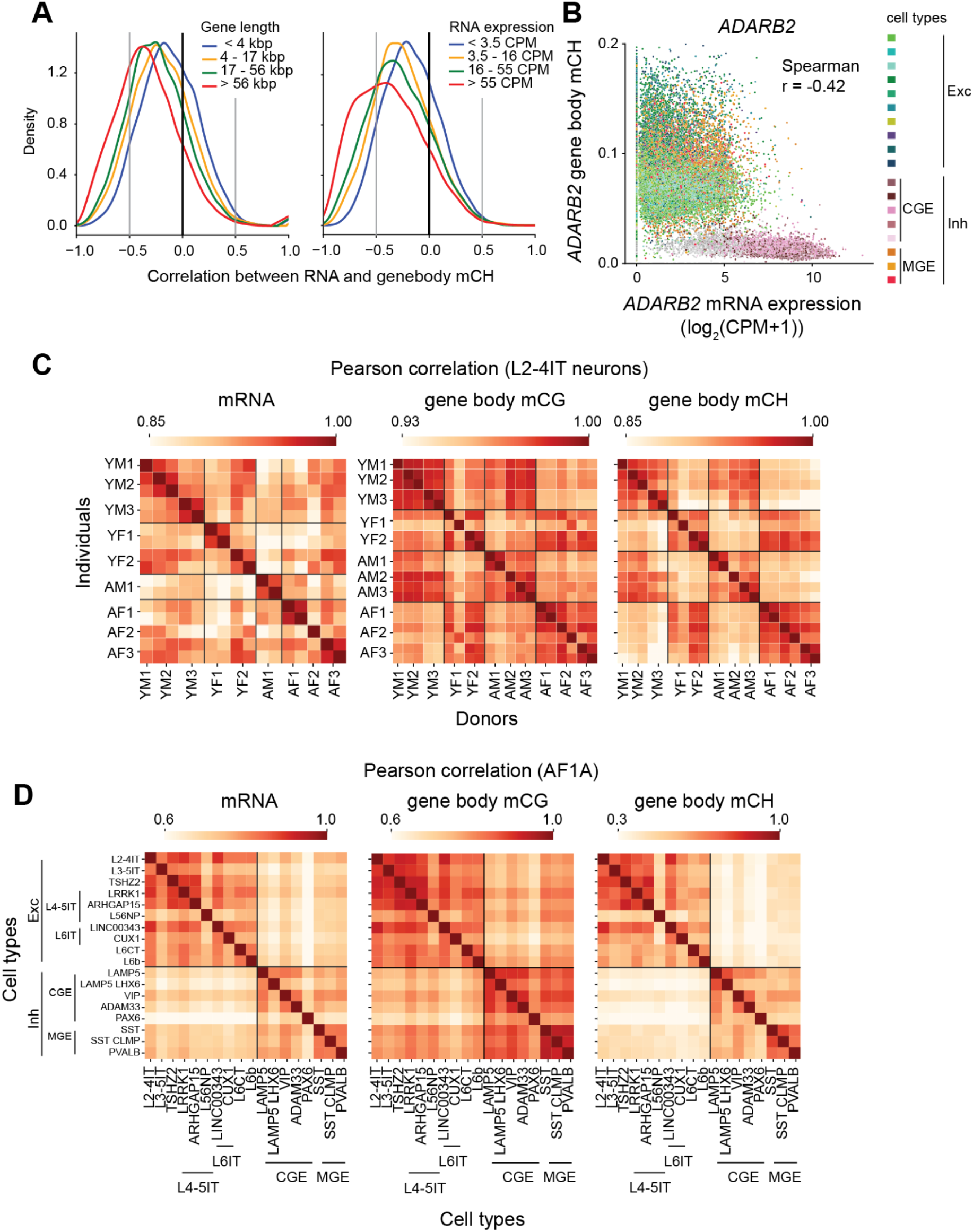
Correlation between samples, cell types and relation between DNA methylation and RNA expression. (A) Spearman correlation between RNA expression and gene body non-CG methylation, stratified by gene length (left) or gene expression (right). (B) The relation between gene body mCH and mRNA expression for each cell colored by cell types. (C) Pearson correlation of mRNA expression (left), gene body CG (middle) and non-CG (right) methylation between samples in L2-4IT neurons. The correlation was calculated by the top 10,000 variation features. (D) Pearson correlation of mRNA expression (left), gene body CG (middle) and non-CG (right) methylation between cell types in one donor. The correlation was calculated by the top 10,000 variation features.

**Fig S4.**
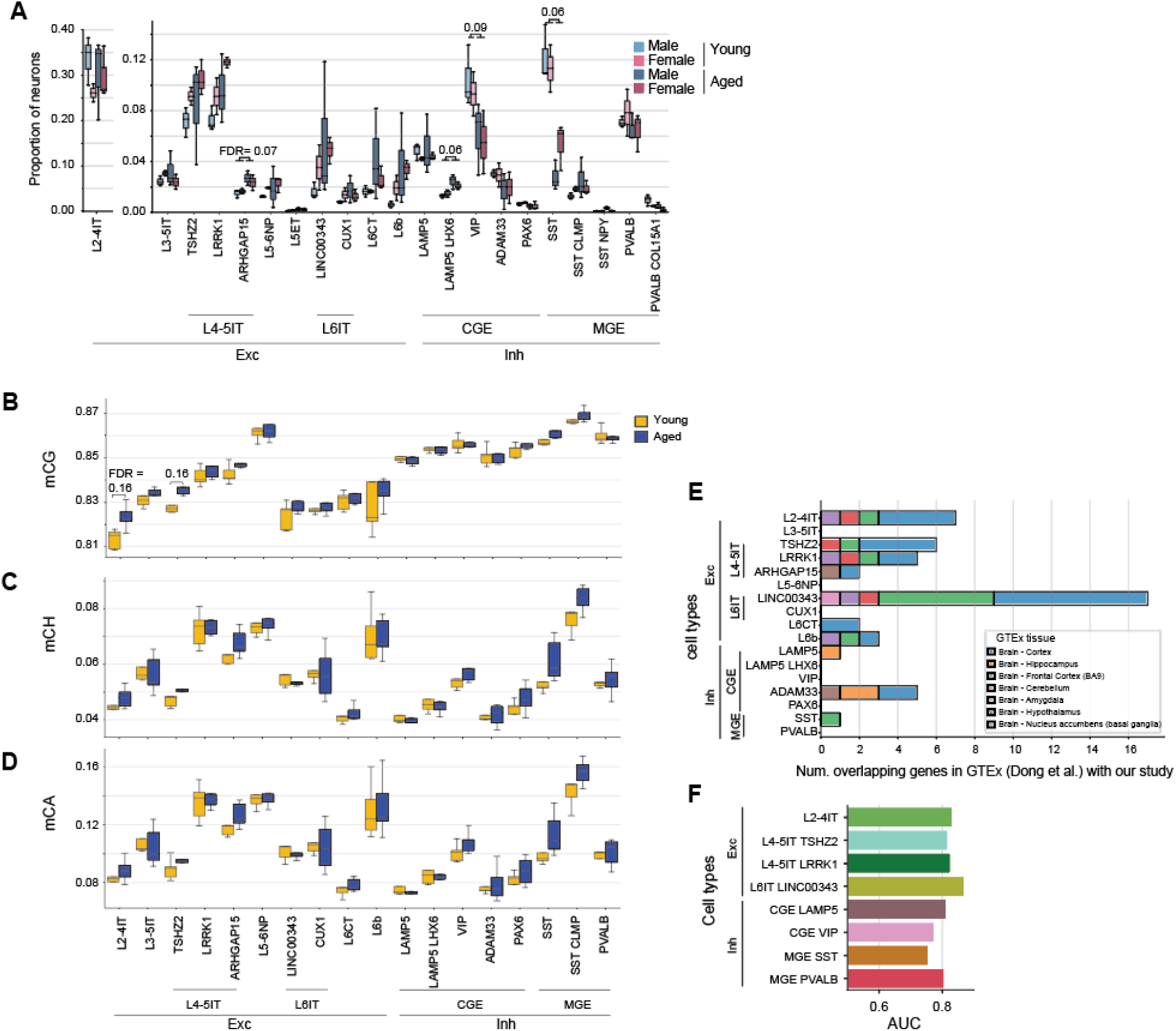
Cell type proportions and global DNA methylation changes in aging. (A) Proportion of all neuron subtypes in young and aged donors. Significant differences between aged and young donor groups are labeled (FDR < 0.1 from Wilcoxon rank-sum test and Benjamini–Hochberg correction, N=5 for young donors, N=6 for aged donors). (B, C, D) Genome wide CG (B), CH (C) and CA (D) methylation in young and aged donors. Significant differences between aged and young donor groups are labeled (FDR < 0.2 from Wilcoxon rank-sum test and Benjamini–Hochberg correction, N=5 for young donors, N=6 for aged donors). (E) Number of age-DEGs in our study overlapping with age-DEGs from GTEx^17^. (F) Cell type prioritization in aging using Augur^22^.

**Fig S5.**
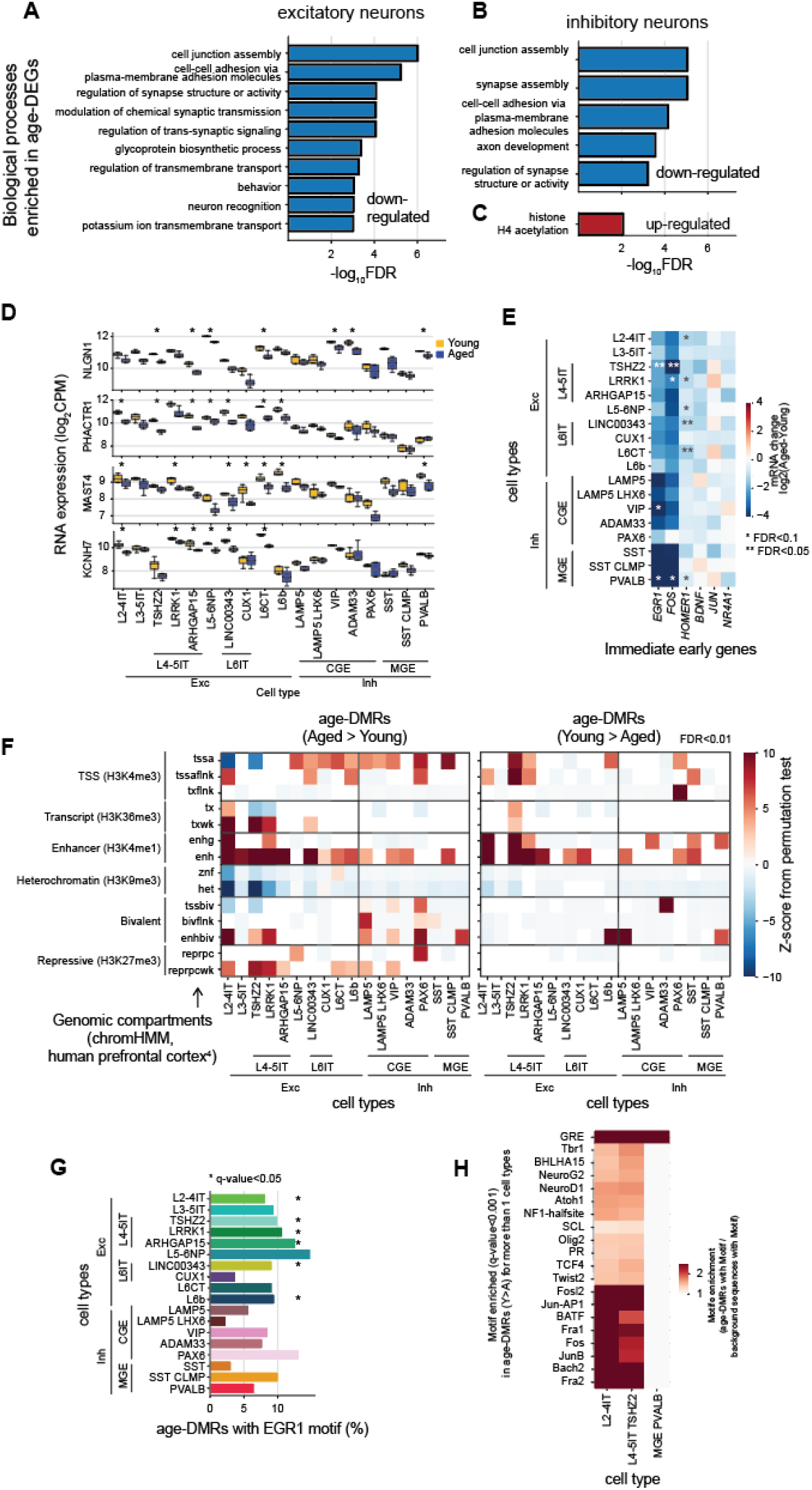
Functional enrichment of age-related transcriptome and DNA methylome change. (A,B) Gene ontology (biological process) enriched (FDR< 0.001) in all down-regulated age-DEGs in (A) excitatory neurons and in (B) inhibitory neurons. (C) Gene ontology (biological process) enriched (FDR< 0.01) all up-regulated age-DEGs in inhibitory neurons. (D) Expression of top 4 genes has significant age effects in most number of cell types in aged and young donors. * significantly age-DEGs (FDR< 0.05 and | fold-change | > 1.2). (E) Log2 fold-change of RNA expression in immediate early genes. *FDR<0.1, **FDR<0.05 (F) Enrichment of age-DMRs in ChromHMM annotated regions. The number of overlapping base pairs was calculated for the set of age-DMRs with each annotation. A baseline was calculated by shuffling the age-DMRs across the genome and overlapping the shuffled DMRs and the annotated regions. The shuffling process was repeated 1,000 times. A normal distribution was fitted for the shuffled results and then compared with the data to get the z-score and p-value. Only significant enrichment/depletion (FDR<0.01) are shown. (G) EGR1 motif enrichment in hypermethylated age-DMRs for all cell types.(*q-value<0.05) (H) Motif enrichment of hypomethylated age-DMRs. Only motifs significantly enriched (q-value<0.001) in more than 1 cell type are shown.

**Fig S6.**
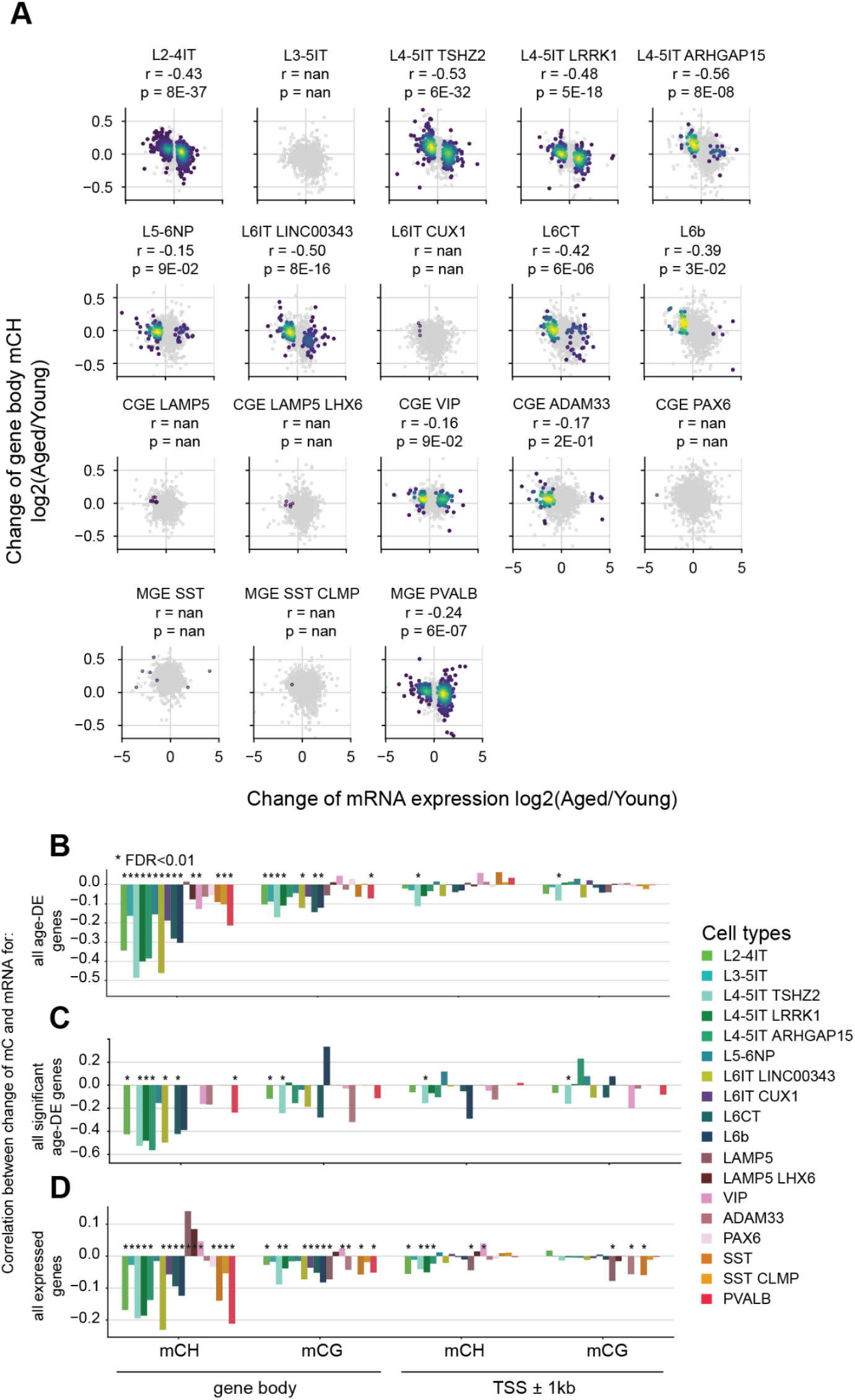
Negative correlation between the change of mRNA expression and DNA methylation in aging. (A) Relationship between age-related change in gene expression (x-axis) and age-related change in gene body mCH (y-axis) for each cell type. Spearman correlation, r. (B-D) Spearman correlation coefficient between the age change of DNA methylation (aged - young) and the fold-change in gene expression in (B) all age-DEGs, (C) significant age-DEGs in that cell type, and (D) all genes. Asterisk marks a significant correlation (FDR <0.01, Benjamini–Hochberg).

**Fig S7.**
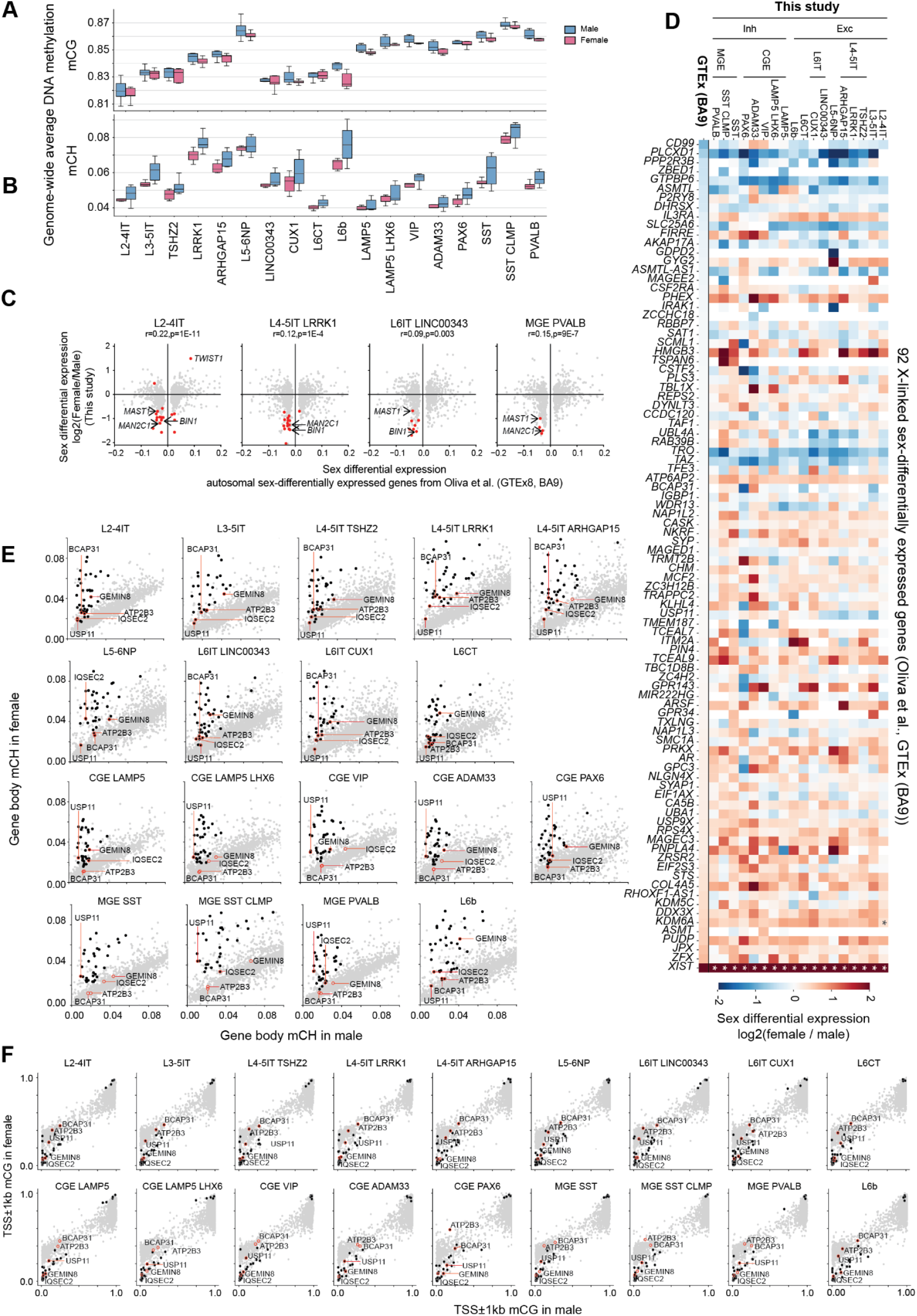
Sex differences in snmCT-seq data (this study) are consistent with published snRNA-seq and WGBS data. (A,B) Genome-wide CG (A) and non-CG (B) methylation in male and female donors. No significant differences between male and female donor groups. (FDR > 0.2 from Wilcoxon rank-sum test and Benjamini–Hochberg correction, N=5 for young donors, N=6 for aged donors). (C) Comparison of the regulation of X-linked RNA expression between females and males in our data and significant age-DEGs in GTEx data ^44^. * significantly age-DEGs in our data (FDR<0.05, |fold-change| > 1.2). (B) Comparison of the regulation of autosomal RNA expression between females and male in our data (y-axis) and significant age-DEGs in GTEx (x-axis). Gray dots: significant age-DEGs in GTEx data. Red dots: Genes with FDR<0.1 and | fold-change | > 1.2 in our data. (E, F) Scatter plots of X-linked gene body mCH (C) or promoter mCG (D) in male and female (gray dots). Black dots are the 41 genes shown in (Fig. 5F). Putative cell type-specific X-inactivation escape genes are labeled.

## Supplementary tables

**Table S1: Donor metadata.** Age, sex, tissue source, and total nuclei passing QC metrics.

**Table S2: Cell type composition.** Number of cells per donor and cell type.

**Table S3: Age differentially expressed genes (age-DEGs).** List of age-DEGs and their statistics analyzed by Dream.

**Table S4: Age differentially methylated regions (age-DMRs).** DMRs called using DSS.

**Table S5: Sex differentially expressed genes (age-DEGs).** List of sex-DEGs and their statistics analyzed by Dream.

**Table S6: Sex differentially methylated regions (sex-DMRs).** DMRs called using DSS.

## METHODS

### Human samples

Human post-mortem brain tissue samples from dorsolateral prefontal cortex (Brodmann area BA46) were acquired via the NIH Neurobiobank and included samples provided by the Brain Tissue Donation Program at the University of Pittsburgh (IRB Protocol Number: REN14120157/IRB981146), University of Miami Brain Endowment Bank (IRB Protocol Number: 19920358 (CR0001775)), and The Human Brain and Spinal Fluid Resource Center (managed by Sepulveda Research Corporation) (IRB Protocol Number: PCC#: 2015-060672, VA Project #: 0002). All donors had no clinical brain diagnosis. For each donor, two independent samples (biological replicates) were obtained either by using separate tissue punches, or in cases where only a single tissue punch was available by separately dissecting cells from opposite sides of a single piece of tissue. RNA integrity (RIN) values were independently tested using an Agilent TapeStation RIN screentape. RNA for the test was isolated from tissue samples using TRIzol-chloroform extraction and eluted on Qiagen RNeasy silica membrane spin columns after DNase I digestion.

### Nuclei isolation, FANS sorting and snmCT sequencing

Nuclei isolation and FANS sorting was performed as previously described ^8,57^, with added protease inhibitors to the homogenization buffer. A detailed protocol can be found at https://www.protocols.io/view/snmcat-v2-x54v9jby1g3e/v2. snmCT-seq library preparation and sequencing was performed as described in ^8,9,58^ and protocol is listed at https://www.protocols.io/view/methyl-c-sequencing-of-single-cell-nuclei-snmc-seq-pjvdkn6

Briefly, postmortem brain chunks were ground in liquid nitrogen and stored at - 80 ℃. The ground tissue was then lysed and fractionated to collect nuclei. The purified single-nuclei were stained with nuclei stain Hoechst 33342 and anti-NeuN antibody AlexaFluor 488 followed by fluorescence-activated nuclei (FANS) sorting into 384-well plates, where cDNA was reverse transcribed and amplified. The samples were bisulfite converted in each well. Finally, the libraries were sequenced by the Illumina NovaSeq 6000 instrument with S4 flowcells and

150-bp paired-end mode.

### Mapping snmCT-seq data

We pre-processed snmCT-seq data as previously described^8,16^ using pipeline (http://cemba-data.readthedocs.io/). The snmCT-seq library contains both DNA- and RNA-derived (cDNA) fragments. To separate these reads from each modality, we mapped reads in both DNA mode and RNA mode and then separated them by the total methylation level in each read. The main steps were: (1) trimming adapters, (2) DNA mapping, (3) call methylation level and extract DNA reads, (4) RNA mapping, (5) call methylation level per read and extract RNA reads, and (6) RNA expression quantification.

1. Trimming: Cutadapt (v4.1) in pair-end mode was used to trim reads with R1 adapter (AGATCGGAAGAGCACACGTCTGAAC), R2 adapter (AGATCGGAAGAGCGTCGTGTAGGGA), template switching oligo (TSO) (AAGCAGTGGTATCAACGCAGAGTGAATGG), N6 (AAGCAGTGGTATCAACGCAGAGTAC), ISPCR(AAGCAGTGGTATCAACGCAGAGT), poly T and polyA. 10 bps from both the 3’ and 5’ end are further trimmed. Reads shorter than 30 bps were discarded.
2. DNA mapping: Reads were mapped to the human genome assembly GRCh38 (hg38) using Hisat-3n (v2.2.1-3n-0.0.3) with parameters: `--base-change C,T; --no-spliced-alignment; --no-temp-splicesite; --directional-mapping-reverse`. Unmapped reads (MAPQ=0) were discarded while unique-mapped reads (MAPQ=60) and multi-mapped reads (MAPQ=1) were kept separately. PCR duplicated reads were removed using Picard MarkDuplicates (v2.27.4).
3. Call methylation level and extract DNA reads: Reads with global non-CG methylation level less than 0.5 were collected as DNA reads.
4. RNA mapping: Reads were mapped to hg38 using Hisat2 (v2.2.1) using default parameters. Multi-mapped reads were removed.
5. Call methylation level and extract RNA reads: Reads with global non-CG methylation level greater than 0.9 were collected as RNA reads.
6. RNA expression quantification: featureCounts (v2.0.1) was used to assign reads to gencode v37 genes including both exon and introns (`-t gene`) with fraction setting (`--fraction`) to evenly distributed reads mapped to overlapped features.

### Personalized genome sequencing and mapping

The genomic DNA was extracted from ground frozen postmortem brain tissue. The library preparation and sequencing were performed as described in ^57^.

Reads from personalized genome sequencing were mapped to human genome assembly GRCh38 (hg38) as follows. Reads were first trimmed with Illumina adaptor sequencing using trim_galore (v0.6.7). 10 bp from the 3’ end from read1 and read2 were also removed. The trimmed reads were mapped by bwa mem (v0.7.17), deduplicated by Picard MarkDuplicates (v2.27.4), and SNPs were called using the Genome Analysis Toolkit (GATK, v4.2.6.1) as instructed in the GATK tutorial ^59^. For each donor, cytosines and adjacent downstream(3’) sites which overlap homozygous SNPs called from any donor, were removed for the downstream methylation analysis.

### Cell filtering

The criteria for a cell to pass DNA methylome quality control were: (1) DNA unique mapped read count > 300,000, (2) mapping rate > 0.5, (3) global mCG level > 0.5, (4) global mCH level < 0.2,and (5) global mCCC level < 0.05. The criteria for a cell to pass RNA quality control were: (1) number of genes detected > 500, (2) percent of mitochondria reads < 10%, and (3) doublet score (called by scrublet^60^) < 0.3.

### Feature filtering

100kb methylation bin features were filtered by coverage and blacklist regions. Bins with mean number of covered cytosine <500 or >2000, or overlapped with ENCODE blacklist^61^ regions were removed for the clustering. Genes expressed in fewer than 100 cells were removed for further analysis.

### Clustering and cell type annotation

Clustering was performed using non-CG DNA methylation features in 100kb bins throughout the genome, as previously described^8,9^. The top 20,000 most variable principle components of the 100kb bins were used to construct the KNN graph using scanpy.pp.neighbors with k=25. Next, the KNN graph was used to perform clustering using the Leiden community detection algorithm^62^ with leiden resolution 1. Consensus clustering was performed by repeating the cluster analysis 500 times with the same resolution (0.7) and different random seed. Clusters were then merged using a balanced random forest classifier (imblearn.ensemble.BalancedRandomForestClassifier^63^) trained using 50% of cells per cluster, while limiting the maximum number of cells to 500 in each cluster to avoid large clusters dominating the training dataset. The merging process stops when it reaches the target accuracy (0.96). Overall, the clustering process is done by ConsensusClustering function in ALLCools (https://lhqing.github.io/ALLCools). Clusters with a high duplication ratio or containing less than 300 cells were excluded from the downstream analysis. Clusters were then annotated based on cell type marker gene expression and gene body methylation features.

### Differential expression and gene ontology analysis

We merged nuclei from each sample for each cell type to get peudobulk counts. Protein coding and lncRNA genes from chromosomes 1-22 and X with counts per million (CPM) >5 in at least 2 samples are defined as “expressed genes”. Differential expression testing was performed separately for each cell type and each comparison (aged vs, young or female vs. male) using Dream^64^ (variancePartition v1.28.9, edgeR v3.40.2). Specifically, `calcNormFactors` was used to control for the raw library size from the raw counts. Then, `voomWithDreamWeights` and `dream` were used to prepare for the linear mixed model fit from the normalized count and design matrix. The model included fixed effects of age and sex, and a random intercept by donor: *expression* ∼ *age* + *sex* + (1| *donor*). Finally, after testing the generalized linear mixed model for each gene, genes with absolute fold-change greater than 1.2 and adjusted p-value < 0.05 are defined as differentially expressed genes (DEGs) in our analysis.

To control for the number of nuclei for each cell type from each condition, we also applied the differential expression test after randomly selecting 50 cells from each sample for each major cell type. The subsampling and DEG testing was repeated five times. The median, 25th and 75th quantile of the number of differentially expressed genes are shown.

Functional gene ontology (GO) enrichment analysis was performed using clusterProfiler (v 3.14.3). All expressed genes in each cell type were included as the background gene list. Gene ontology with gene size between 25 and 500 were included in the enrichment test. The Benjamini-Hochberg method was used to control the false discovery rate.

### Differential methylation analysis

We created pseudobulk counts for nuclei from each donor for each cell type and tested each CpG site between each condition comparison (aged vs. young or female vs. male) to detect differentially methylated regions (DMRs) using DSS^20,21^ (DSS v2.34.0, bsseq v1.22.0). `DMLfit.multiFactor` was applied with model formula: *mCG*∼*age* + *sex*, with smoothing window size 500 bp. Next, `DMLtest.multiFactor` was used to call differentially methylated loci. Finally, `callDMR` was used to merge significantly differentially methylated loci (p<0.05) with a distance smaller than 50 bp. Merged regions with a length greater than 50 bp and containing at least 3 CpG sites are called differentially methylated regions (DMRs).

To control for the number of nuclei for each cell type from each condition, we also apply the differential methylation test after randomly subsampling 100 cells from each donor for major cell types. The subsampling was done five times. The median, 25th and 75th quantile of the number of differentially methylated regions are shown.

### Differentially methylated region (DMR) enrichment test

We utilized the chromHMM annotations of 15 chromatin states derived from human frontal cortex tissue^29^. For each chromatin state, we computed the overlap in base pairs (Ntrue) between the DMRs and the designated state regions. Subsequently, this process was repeated 1,000 times, each time with the DMRs randomly shuffled across the genome. The number of base pairs overlapping between shuffled DMRs and the chromatin state region are fitted using a normal distribution. Finally, Ntrue was used to compare with the distribution to derive both the z-score and the associated p-value. The Benjamini-Hochberg method was used to control false discovery rate.

### DNA methylation valleys (DMVs)

To cell DNA methylation valleys (DMVs), we first identified undermethylated regions (UMRs) and low methylated regions (LMRs) using MethylSeekR^65^. We used methylation cutoff m=0.5, minimum number of CpG, n=10, and FDR < 0.05 as parameters for detecting UMRs and LMRs. DMVs were defined as UMRs with length ≥5 kb and mean methylation level ≤15%. We detected 645-1546 DMVs in each cell type and 35%-63% of them overlapped with the DMVs detected in human embryonic stem cell-derived neural progenitor cells^35^.

### Estimation of telomere content from snmCT-seq

We estimated telomere content as a proxy of telomere length from snmCT-seq dataset using TelomereHunter (v1.1.0) ^40^. In short, TelomereHunter first searched for reads with telomeric repeats from pre-aligned BAM files, and then classified the selected reads into four categories: intratelomeric, junction spanning, subtelomeric and intrachromosomal, using their alignment information.

We first generated per-cell BAM files that contained unmapped reads by re-running Hisat-3n, followed by de-duplication using Picard MarkDuplicates. Next, RNA reads identified from our previous steps described in the “Mapping snmCT-seq data” section were excluded. The resulting BAM files were used as the input of Telomerehunter to search for t-type (TTAGGG) telomeric repeats as well as c-type (TCAGGG), g-type (TGAGGG) and j-type (TTGGGG) telomeric variant repeats. Reads were considered as telomeric reads if they contained at least 6 repeats per 100 bp read length. Given the number of reads of comparable GC content (48-52%) was low per cell, we used the number of uniquely mapped DNA reads in each snmCT-seq library as the normalization factor. The telomere content for each cell was computed as the number of intratelomeric reads divided by the number of uniquely mapped DNA reads, and multiplied by 10^6^.

## References

1. BRAIN Initiative Cell Census Network (BICCN) (2021). A multimodal cell census and atlas of the mammalian primary motor cortex. Nature 598, 86–102.

2. Johansen, N., Somasundaram, S., Travaglini, K.J., Yanny, A.M., Shumyatcher, M., Casper, T., Cobbs, C., Dee, N., Ellenbogen, R., Ferreira, M., et al. (2023). Interindividual variation in human cortical cell type abundance and expression. Science 382, eadf2359.

3. Lister, R., Mukamel, E.A., Nery, J.R., Urich, M., Puddifoot, C.A., Johnson, N.D., Lucero, J., Huang, Y., Dwork, A.J., Schultz, M.D., et al. (2013). Global epigenomic reconfiguration during mammalian brain development. Science 341, 1237905.

4. Jaffe, A.E., Gao, Y., Deep-Soboslay, A., Tao, R., Hyde, T.M., Weinberger, D.R., and Kleinman, J.E. (2016). Mapping DNA methylation across development, genotype and schizophrenia in the human frontal cortex. Nat. Neurosci. 19, 40–47.

5. Herring, C.A., Simmons, R.K., Freytag, S., Poppe, D., Moffet, J.J.D., Pflueger, J., Buckberry, S., Vargas-Landin, D.B., Clément, O., Echeverría, E.G., et al. (2022). Human prefrontal cortex gene regulatory dynamics from gestation to adulthood at single-cell resolution. Cell 185, 4428–4447.e28.

6. Li, J., Jaiswal, M.K., Chien, J.-F., Kozlenkov, A., Jung, J., Zhou, P., Gardashli, M., Pregent, L.J., Engelberg-Cook, E., Dickson, D.W., et al. (2023). Divergent single cell transcriptome and epigenome alterations in ALS and FTD patients with C9orf72 mutation. Nat. Commun. 14, 1–22.

7. Mathys, H., Peng, Z., Boix, C.A., Victor, M.B., Leary, N., Babu, S., Abdelhady, G., Jiang, X., Ng, A.P., Ghafari, K., et al. (2023). Single-cell atlas reveals correlates of high cognitive function, dementia, and resilience to Alzheimer’s disease pathology. Cell 186, 4365–4385.e27.

8. Luo, C., Liu, H., Xie, F., Armand, E.J., Siletti, K., and Bakken, T.E. (2022). Single nucleus multi-omics identifies human cortical cell regulatory genome diversity. Cell Genomics.

9. Luo, C., Keown, C.L., Kurihara, L., Zhou, J., He, Y., Li, J., Castanon, R., Lucero, J., Nery, J.R., Sandoval, J.P., et al. (2017). Single-cell methylomes identify neuronal subtypes and regulatory elements in mammalian cortex. Science 357, 600–604.

10. Hodge, R.D., Bakken, T.E., Miller, J.A., Smith, K.A., Barkan, E.R., Graybuck, L.T., Close, J.L., Long, B., Johansen, N., Penn, O., et al. (2019). Conserved cell types with divergent features in human versus mouse cortex. Nature 573, 61–68.

11. Lauterborn, J.C., Scaduto, P., Cox, C.D., Schulmann, A., Lynch, G., Gall, C.M., Keene, C.D., and Limon, A. (2021). Increased excitatory to inhibitory synaptic ratio in parietal cortex samples from individuals with Alzheimer’s disease. Nat. Commun. 12, 2603.

12. Song, Y.-H., Yoon, J., and Lee, S.-H. (2021). The role of neuropeptide somatostatin in the brain and its application in treating neurological disorders. Exp. Mol. Med. 53, 328–338.

13. Rozycka, A., and Liguz-Lecznar, M. (2017). The space where aging acts: focus on the GABAergic synapse. Aging Cell 16, 634–643.

14. Gabitto, M., Travaglini, K., Ariza, J., Kaplan, E., Long, B., Rachleff, V., Ding, Y., Mahoney, J., Dee, N., Goldy, J., et al. (2023). Integrated multimodal cell atlas of Alzheimer’s disease. Res Sq. 10.21203/rs.3.rs-2921860/v1.

15. Kozlenkov, A., Li, J., Apontes, P., Hurd, Y.L., Byne, W.M., Koonin, E.V., Wegner, M., Mukamel, E.A., and Dracheva, S. (2018). A unique role for DNA (hydroxy)methylation in epigenetic regulation of human inhibitory neurons. Science Advances 4, eaau6190.

16. Liu, H., Zhou, J., Tian, W., Luo, C., Bartlett, A., Aldridge, A., Lucero, J., Osteen, J.K., Nery, J.R., Chen, H., et al. (2021). DNA methylation atlas of the mouse brain at single-cell resolution. Nature 598, 120–128.

17. Dong, X., Sun, S., Zhang, L., Kim, S., Tu, Z., Montagna, C., Maslov, A.Y., Suh, Y., Wang, T., Campisi, J., et al. (2021). Age-related telomere attrition causes aberrant gene expression in sub-telomeric regions. Aging Cell 20, e13357.

18. Horvath, S. (2013). DNA methylation age of human tissues and cell types. Genome Biol. 14, R115.

19. Johnstone, S.E., Gladyshev, V.N., Aryee, M.J., and Bernstein, B.E. (2022). Epigenetic clocks, aging, and cancer. Science 378, 1276–1277.

20. Feng, H., Conneely, K.N., and Wu, H. (2014). A Bayesian hierarchical model to detect differentially methylated loci from single nucleotide resolution sequencing data. Nucleic Acids Res. 42, e69.

21. Park, Y., and Wu, H. (2016). Differential methylation analysis for BS-seq data under general experimental design. Bioinformatics 32, 1446–1453.

22. Skinnider, M.A., Squair, J.W., Kathe, C., Anderson, M.A., Gautier, M., Matson, K.J.E., Milano, M., Hutson, T.H., Barraud, Q., Phillips, A.A., et al. (2020). Cell type prioritization in single-cell data. Nat. Biotechnol. 10.1038/s41587-020-0605-1.

23. Kwon, H.-B., Kozorovitskiy, Y., Oh, W.-J., Peixoto, R.T., Akhtar, N., Saulnier, J.L., Gu, C., and Sabatini, B.L. (2012). Neuroligin-1-dependent competition regulates cortical synaptogenesis and synapse number. Nat. Neurosci. 15, 1667–1674.

24. Hamada, N., Ogaya, S., Nakashima, M., Nishijo, T., Sugawara, Y., Iwamoto, I., Ito, H., Maki, Y., Shirai, K., Baba, S., et al. (2018). De novo PHACTR1 mutations in West syndrome and their pathophysiological effects. Brain 141, 3098–3114.

25. Shi, W., Wymore, R.S., Wang, H.S., Pan, Z., Cohen, I.S., McKinnon, D., and Dixon, J.E. (1997). Identification of two nervous system-specific members of the erg potassium channel gene family. J. Neurosci. 17, 9423–9432.

26. Lu, T., Pan, Y., Kao, S.-Y., Li, C., Kohane, I., Chan, J., and Yankner, B.A. (2004). Gene regulation and DNA damage in the ageing human brain. Nature 429, 883–891.

27. Kozlova, I., Sah, S., Keable, R., Leshchyns’ka, I., Janitz, M., and Sytnyk, V. (2020). Cell Adhesion Molecules and Protein Synthesis Regulation in Neurons. Front. Mol. Neurosci. 13, 592126.

28. Mo, A., Mukamel, E.A., Davis, F.P., Luo, C., Henry, G.L., Picard, S., Urich, M.A., Nery, J.R., Sejnowski, T.J., Lister, R., et al. (2015). Epigenomic Signatures of Neuronal Diversity in the Mammalian Brain. Neuron 86, 1369–1384.

29. Roadmap Epigenomics Consortium, Kundaje, A., Meuleman, W., Ernst, J., Bilenky, M., Yen, A., Heravi-Moussavi, A., Kheradpour, P., Zhang, Z., Wang, J., et al. (2015). Integrative analysis of 111 reference human epigenomes. Nature 518, 317–330.

30. Heinz, S., Benner, C., Spann, N., Bertolino, E., Lin, Y.C., Laslo, P., Cheng, J.X., Murre, C., Singh, H., and Glass, C.K. (2010). Simple combinations of lineage-determining transcription factors prime cis-regulatory elements required for macrophage and B cell identities. Mol. Cell 38, 576–589.

31. Duclot, F., and Kabbaj, M. (2017). The Role of Early Growth Response 1 (EGR1) in Brain Plasticity and Neuropsychiatric Disorders. Front. Behav. Neurosci. 11, 35.

32. Jones, M.W., Errington, M.L., French, P.J., Fine, A., Bliss, T.V., Garel, S., Charnay, P., Bozon, B., Laroche, S., and Davis, S. (2001). A requirement for the immediate early gene Zif268 in the expression of late LTP and long-term memories. Nat. Neurosci. 4, 289–296.

33. Penner, M.R., Parrish, R.R., Hoang, L.T., Roth, T.L., Lubin, F.D., and Barnes, C.A. (2016). Age-related changes in Egr1 transcription and DNA methylation within the hippocampus. Hippocampus 26, 1008–1020.

34. Hannum, G., Guinney, J., Zhao, L., Zhang, L., Hughes, G., Sadda, S., Klotzle, B., Bibikova, M., Fan, J.-B., Gao, Y., et al. (2013). Genome-wide methylation profiles reveal quantitative views of human aging rates. Mol. Cell 49, 359–367.

35. Xie, W., Schultz, M.D., Lister, R., Hou, Z., Rajagopal, N., Ray, P., Whitaker, J.W., Tian, S., Hawkins, R.D., Leung, D., et al. (2013). Epigenomic analysis of multilineage differentiation of human embryonic stem cells. Cell 153, 1134–1148.

36. Zhang, Y., Amaral, M.L., Zhu, C., Grieco, S.F., Hou, X., Lin, L., Buchanan, J., Tong, L., Preissl, S., Xu, X., et al. (2022). Single-cell epigenome analysis reveals age-associated decay of heterochromatin domains in excitatory neurons in the mouse brain. Cell Res. 10.1038/s41422-022-00719-6.

37. Zannas, A.S., Jia, M., Hafner, K., Baumert, J., Wiechmann, T., Pape, J.C., Arloth, J., Ködel, M., Martinelli, S., Roitman, M., et al. (2019). Epigenetic upregulation of FKBP5 by aging and stress contributes to NF-κB-driven inflammation and cardiovascular risk. Proc. Natl. Acad. Sci. U. S. A. 116, 11370–11379.

38. Blair, L.J., Nordhues, B.A., Hill, S.E., Scaglione, K.M., O’Leary, J.C., 3rd, Fontaine, S.N., Breydo, L., Zhang, B., Li, P., Wang, L., et al. (2013). Accelerated neurodegeneration through chaperone-mediated oligomerization of tau. J. Clin. Invest. 123, 4158–4169.

39. Fullerton, E.F., Karom, M.C., Streicher, J.M., Young, L.J., and Murphy, A.Z. (2022). Age-Induced Changes in μ-Opioid Receptor Signaling in the Midbrain Periaqueductal Gray of Male and Female Rats. J. Neurosci. 42, 6232–6242.

40. Feuerbach, L., Sieverling, L., Deeg, K.I., Ginsbach, P., Hutter, B., Buchhalter, I., Northcott, P.A., Mughal, S.S., Chudasama, P., Glimm, H., et al. (2019). TelomereHunter - in silico estimation of telomere content and composition from cancer genomes. BMC Bioinformatics 20, 272.

41. López-Otín, C., Blasco, M.A., Partridge, L., Serrano, M., and Kroemer, G. (2013). The hallmarks of aging. Cell 153, 1194–1217.

42. Di Micco, R., Krizhanovsky, V., Baker, D., and d’Adda di Fagagna, F. (2021). Cellular senescence in ageing: from mechanisms to therapeutic opportunities. Nat. Rev. Mol. Cell Biol. 22, 75–95.

43. Robin, J.D., Ludlow, A.T., Batten, K., Magdinier, F., Stadler, G., Wagner, K.R., Shay, J.W., and Wright, W.E. (2014). Telomere position effect: regulation of gene expression with progressive telomere shortening over long distances. Genes Dev. 28, 2464–2476.

44. Oliva, M., Muñoz-Aguirre, M., Kim-Hellmuth, S., Wucher, V., Gewirtz, A.D.H., Cotter, D.J., Parsana, P., Kasela, S., Balliu, B., Viñuela, A., et al. (2020). The impact of sex on gene expression across human tissues. Science 369. 10.1126/science.aba3066.

45. Keown, C.L., Berletch, J.B., Castanon, R., Nery, J.R., Disteche, C.M., Ecker, J.R., and Mukamel, E.A. (2017). Allele-specific non-CG DNA methylation marks domains of active chromatin in female mouse brain. Proc. Natl. Acad. Sci. U. S. A. 114, E2882–E2890.

46. Schultz, M.D., He, Y., Whitaker, J.W., Hariharan, M., Mukamel, E.A., Leung, D., Rajagopal, N., Nery, J.R., Urich, M.A., Chen, H., et al. (2015). Human body epigenome maps reveal noncanonical DNA methylation variation. Nature 523, 212–216.

47. Lopes-Ramos, C.M., Chen, C.-Y., Kuijjer, M.L., Paulson, J.N., Sonawane, A.R., Fagny, M., Platig, J., Glass, K., Quackenbush, J., and DeMeo, D.L. (2020). Sex Differences in Gene Expression and Regulatory Networks across 29 Human Tissues. Cell Rep. 31, 107795.

48. Fu, X.-M., and Zhu, B.T. (2009). Human pancreas-specific protein disulfide isomerase homolog (PDIp) is an intracellular estrogen-binding protein that modulates estrogen levels and actions in target cells. J. Steroid Biochem. Mol. Biol. 115, 20–29.

49. Gusev, F.E., Reshetov, D.A., Mitchell, A.C., Andreeva, T.V., Dincer, A., Grigorenko, A.P., Fedonin, G., Halene, T., Aliseychik, M., Filippova, E., et al. (2019). Chromatin profiling of cortical neurons identifies individual epigenetic signatures in schizophrenia. Transl. Psychiatry 9, 256.

50. Tukiainen, T., Villani, A.-C., Yen, A., Rivas, M.A., Marshall, J.L., Satija, R., Aguirre, M., Gauthier, L., Fleharty, M., Kirby, A., et al. (2017). Landscape of X chromosome inactivation across human tissues. Nature 550, 244–248.

51. Haghani, A., Li, C.Z., Robeck, T.R., Zhang, J., Lu, A.T., Ablaeva, J., Acosta-Rodríguez, V.A., Adams, D.M., Alagaili, A.N., Almunia, J., et al. (2023). DNA methylation networks underlying mammalian traits. Science 381, eabq5693.

52. Dileep, V., Boix, C.A., Mathys, H., Marco, A., Welch, G.M., Meharena, H.S., Loon, A., Jeloka, R., Peng, Z., Bennett, D.A., et al. (2023). Neuronal DNA double-strand breaks lead to genome structural variations and 3D genome disruption in neurodegeneration. Cell 186, 4404–4421.e20.

53. Ain, Q., Schmeer, C., Penndorf, D., Fischer, M., Bondeva, T., Förster, M., Haenold, R., Witte, O.W., and Kretz, A. (2018). Cell cycle-dependent and -independent telomere shortening accompanies murine brain aging. Aging 10, 3397–3420.

54. Bakken, T.E., Jorstad, N.L., Hu, Q., Lake, B.B., Tian, W., Kalmbach, B.E., Crow, M., Hodge, R.D., Krienen, F.M., Sorensen, S.A., et al. (2021). Comparative cellular analysis of motor cortex in human, marmoset and mouse. Nature 598, 111–119.

55. Tasic, B., Yao, Z., Graybuck, L.T., Smith, K.A., Nguyen, T.N., Bertagnolli, D., Goldy, J., Garren, E., Economo, M.N., Viswanathan, S., et al. (2018). Shared and distinct transcriptomic cell types across neocortical areas. Nature 563, 72–78.

56. Allen, W.E., Blosser, T.R., Sullivan, Z.A., Dulac, C., and Zhuang, X. (2023). Molecular and spatial signatures of mouse brain aging at single-cell resolution. Cell 186, 194–208.e18.

57. Tian, W., Zhou, J., Bartlett, A., Zeng, Q., Liu, H., Castanon, R.G., Kenworthy, M., Altshul, J., Valadon, C., Aldridge, A., et al. (2023). Single-cell DNA methylation and 3D genome architecture in the human brain. Science 382, eadf5357.

58. Luo, C., Rivkin, A., Zhou, J., Sandoval, J.P., Kurihara, L., Lucero, J., Castanon, R., Nery, J.R., Pinto-Duarte, A., Bui, B., et al. (2018). Robust single-cell DNA methylome profiling with snmC-seq2. Nat. Commun. 9, 3824.

59. Van der Auwera, G.A., and O’Connor, B.D. (2020). Genomics in the Cloud: Using Docker, GATK, and WDL in Terra (“O’Reilly Media, Inc.”).

60. Wolock, S.L., Lopez, R., and Klein, A.M. (2019). Scrublet: Computational Identification of Cell Doublets in Single-Cell Transcriptomic Data. Cell Syst 8, 281–291.e9.

61. Amemiya, H.M., Kundaje, A., and Boyle, A.P. (2019). The ENCODE Blacklist: Identification of Problematic Regions of the Genome. Sci. Rep. 9, 9354.

62. Traag, V.A., Waltman, L., and van Eck, N.J. (2019). From Louvain to Leiden: guaranteeing well-connected communities. Sci. Rep. 9, 5233.

63. Lemaître, G., Nogueira, F., and Aridas, C.K. (2017). Imbalanced-learn: A Python Toolbox to Tackle the Curse of Imbalanced Datasets in Machine Learning. J. Mach. Learn. Res. 18, 1–5.

64. Hoffman, G.E., and Roussos, P. (2021). Dream: powerful differential expression analysis for repeated measures designs. Bioinformatics 37, 192–201.

65. Burger, L., Gaidatzis, D., Schübeler, D., and Stadler, M.B. (2013). Identification of active regulatory regions from DNA methylation data. Nucleic Acids Res. 41, e155.

